# Universal rules govern plasmid copy number

**DOI:** 10.1101/2024.10.04.616648

**Authors:** Paula Ramiro-Martínez, Ignacio de Quinto, Val F. Lanza, João Alves Gama, Jerónimo Rodríguez-Beltrán

## Abstract

Plasmids –autonomously replicating DNA molecules– exhibit a broad range of replication and mobility strategies, genetic repertoires, host ranges, sizes, and copies per cell. However, the determinants of plasmid copy number (PCN) remain poorly understood. Here, we use extensive DNA sequencing data to analyse the copy number of thousands of diverse bacterial plasmids in a comprehensive manner. We find that PCN is highly variable, spanning nearly three orders of magnitude, and that it is intrinsically robust against changes in genomic context. We further show that PCN variability is tightly associated with plasmid lifestyles, and propose the concept of replicon dominance to explain interactions in widespread multi-replicon plasmids. Finally, we uncover a universal scaling law that links copy number and plasmid size across bacterial species, indicating that pervasive constraints modulate the PCN-size trade-off.

## Introduction

Plasmids are typically circular, autonomously replicating DNA molecules, that stably co-exist with host chromosomes. As the main drivers of horizontal gene transfer, plasmids can cross phylogenetic boundaries and be present in different microbial genera, families, and even life domains^1^. Plasmids are pervasive and show a plethora of replication and mobility strategies, lengths, host ranges, topologies, G+C contents, and genetic repertoires, including antibiotic resistance and virulence genes^2–6^.

Plasmids ensure their stability in microbial populations thanks to fine-tuned replication mechanisms that maintain a given number of plasmid copies per cell. Plasmid copy number (PCN) is thus a fundamental aspect of plasmid biology that governs plasmid lifestyles. Small plasmids typically lack active partition systems, so they are randomly distributed (segregated) to daughter cells. To avoid being stochastically lost during cell division, these plasmids rely on being present at a high PCN, which statistically guarantees their stable inheritance and persistence in the population^7^. On the other hand, large plasmids are typically present at low PCN as they carry active segregation and partition systems that mechanistically ensure their persistence. Their low PCN likely reduces their metabolic load to the host, alleviating their fitness cost^8^. Therefore, copy number and size are highly intertwined plasmid properties that have been shown to be negatively correlated^9–11^.

Moreover, PCN modulates plasmid evolvability. A high PCN increases the dosage and, proportionally, the expression of plasmid-encoded genes, which is advantageous under antibiotic pressure or in many other stressful environments^12,13^. In addition, variation in PCN within populations generates heterogeneity in gene expression, facilitating bacterial adaptation through phenotypic plasticity^1,14,15^. At longer timescales, copy number determines the evolution of plasmid genes by affecting key parameters such as mutation and recombination rates or genetic drift^1,16,17^.

Despite its paramount importance for microbial biology and evolution, PCN remains relatively understudied. Traditionally, PCN determinations have relied on burdensome experimental techniques (e.g., qPCR, Southern blot)^18^ and are mainly limited to well-known model plasmids (albeit with some exceptions^9^). In this work, we developed a custom bioinformatic pipeline that leverages extensive DNA sequencing data from different studies to calculate PCN for 6,327 phylogenetically diverse plasmids. Our results show that PCN is highly variable among individual plasmids. Still, each plasmid maintains a characteristic PCN that is generally stable, regardless of other plasmids, and across hosts and genetic cargos. We describe the intrinsic sources of PCN variation across plasmids and uncover the principles driving the PCN of multi-replicon plasmids. Moreover, our results reveal a conserved negative relationship between plasmid size and PCN that is independent of host phylogeny. We discover that independently of the plasmid size or replication type, any given plasmid comprises ∼2.5% of the chromosome size of its host. Altogether, our results provide the first large-scale dataset of PCNs across plasmid groups while uncovering a universal scaling law that governs plasmid biology.

## Results

### A database of complete plasmid sequences and their copy number

To comprehensively understand the driving factors of PCN, we established a database of high-quality closed plasmid sequences found in bacterial genomes belonging to nine different bacterial genera from two distinct phyla: Pseudomonadota and Bacillota (henceforth referred to as Gram-negative and Gram-positive, respectively). The selected genera comprised 95 species with biotechnological and clinical interest, such as all members of the ESKAPEE group^19^ (Figure 1). We extracted plasmid sequences from these genomes and classified them into plasmid groups according to their replication mechanism (replicon types^20^) and similarity across whole plasmid sequence content (plasmid taxonomic units – PTUs^21^, and plasmid clusters). This approach gave rise to a dataset that comprises plasmids belonging to 139 PTUs, 238 distinct replicon types, and 2,200 sequence-based clusters, indicating that it captures a significant fraction of the extant plasmid diversity. To estimate PCN for each of these plasmids, we calculated the trimmed mean sequencing coverage of each plasmid relative to the coverage of their host chromosome (see methods), leading to a dataset comprising 6,327 closed, circular, high-quality plasmid sequences and their respective copy numbers (Figure 1, Supplementary Fig. 1, Supplementary Dataset 1 and Supplementary Dataset 2).

**Figure 1.**
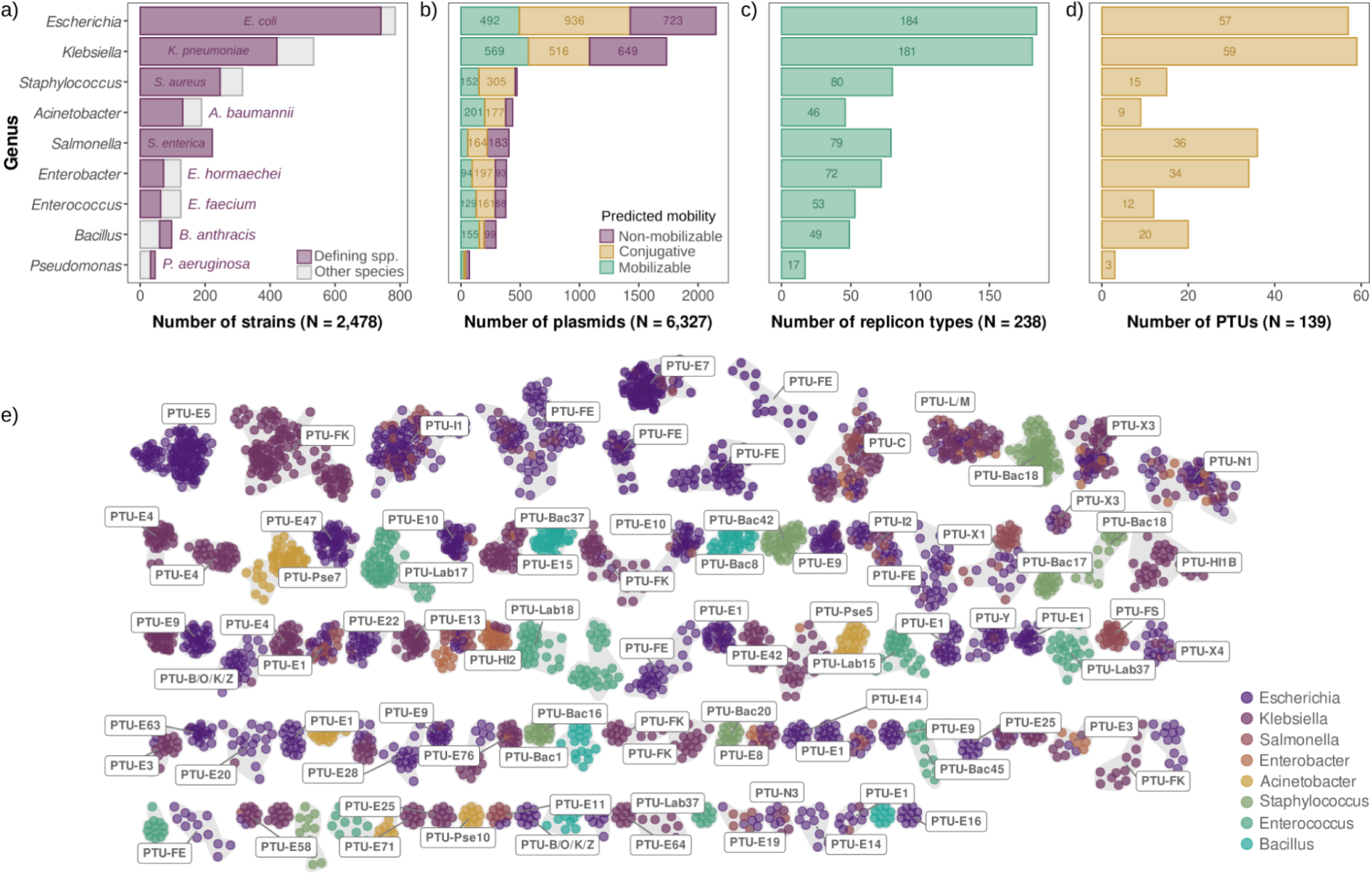
The plasmid dataset. **a)** Number of genomes (x-axis) per genus (y-axis). The most frequent species of each genus is represented in purple, while the rest are grouped and represented in grey. **b)** Number of plasmids (x-axis) per genus classified according to their predicted mobility. **c)** Number of distinct replicon types, and **d)** plasmid taxonomic units (PTUs) (x-axis) per genus. **e)** Plasmid clustering. Each node represents a plasmid, coloured according to the genus in which it was present. Clusters associated with a known PTU are indicated with a label. Clusters with less than ten plasmids are shown in Supplementary Fig. 1.

### PCN is associated with host phylogeny, plasmid mobility, and plasmid groups

PCN was extremely variable in our dataset. In a logarithmic scale, PCNs displayed a broad bimodal distribution spanning three orders of magnitude (Figure 2a), reflecting two well-known plasmid lifestyle strategies^1,2^: Low-copy number plasmids (LCPs), typically ranging from 1 to 2 copies per chromosome (mode = 1.49) and high-copy number plasmids (HCPs), usually bearing more than ten copies per cell (mode = 10.40). Hereafter, we use the anti-mode of this distribution (i.e., the valley between both peaks: 5.75 copies) as a threshold to differentiate LCPs from HCPs. Although this threshold was largely consistent with plasmid sizes (Supplementary Fig. 2) and previous non-empirical definitions^7^, we note that it is likely biased by the overrepresentation of Enterobacterales in our dataset and that the anti-mode in PCN distributions was not equally evident across phylogenetic groups (Figure 2b). Regardless of the shape of the distribution, HCPs and LCPs were present in all genera, although at different proportions: HCPs were more often found in *Escherichia* and *Enterobacter,* while *Pseudomonas*, *Enterococcus*, *Bacillus*, and *Klebsiella* were significantly enriched in LCPs (Chi-squared test, Benjamini-Hochberg (BH) adjusted p < 10^-3^ and Cohen’s h (effect size) > 0.1 in all cases; Figure 2c).

**Figure 2.**
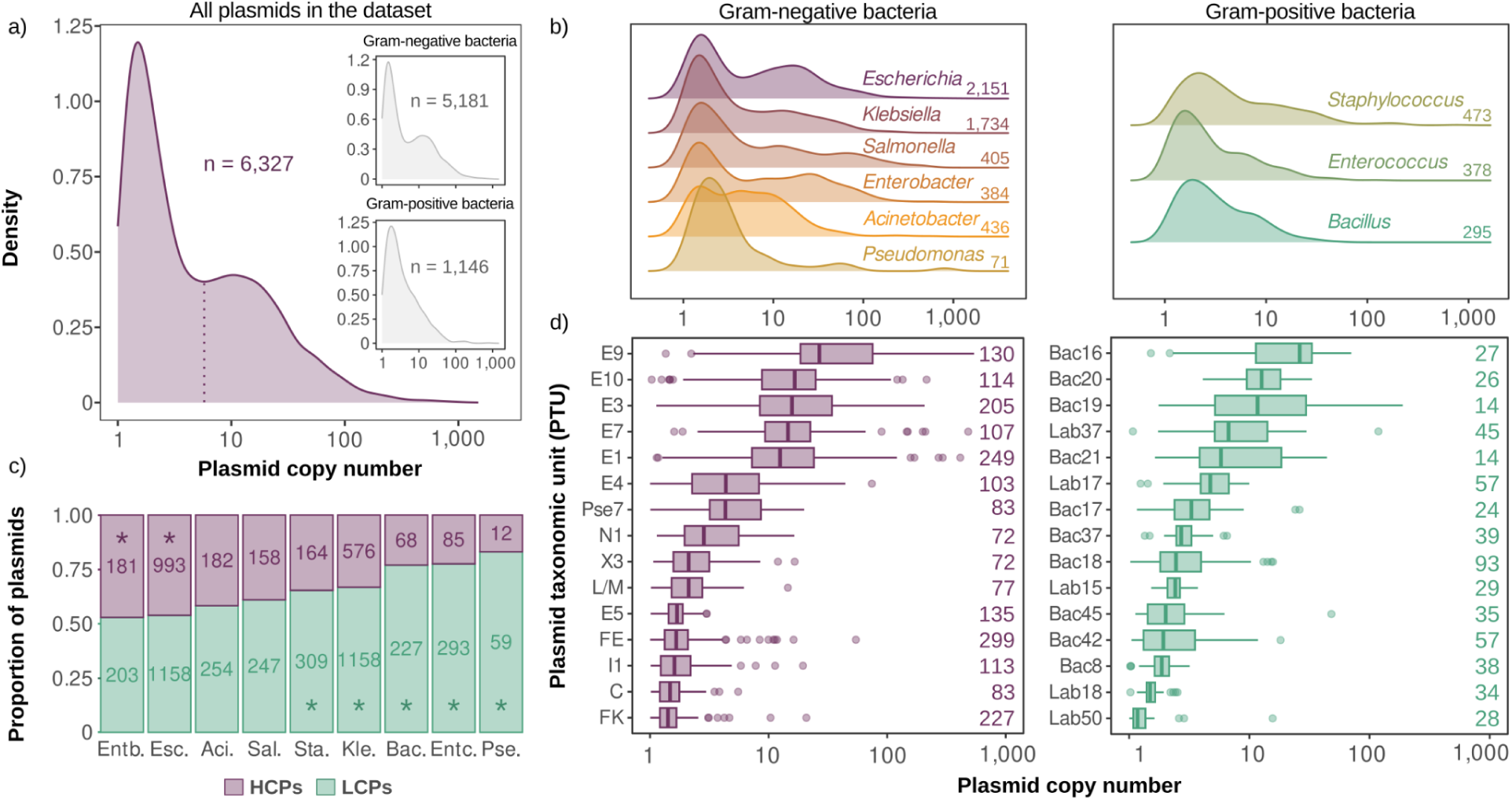
PCN is associated with host phylogeny and plasmid groups. **a)** PCN distribution for all analysed plasmids (n = 6,327). Inset plots represent the same plasmids separated according to the classification of their hosts. The dotted line represents the anti-mode (5.75 copies) of the distribution. **b)** Distribution of PCN within each host genus. The numbers on the right represent the number of plasmids for each genus. **c)** Proportion of HCPs (purple) and LCPs (green) (y-axis) by host genus (x-axis). The numbers within each bar denote the number of plasmids belonging to each category. Asterisks denote that the group of plasmids where they are placed (HCPs or LCPs) is significantly overrepresented compared to the complete plasmid dataset (white dashed line). The x-axis represents host genera, abbreviated to three (or four) letters. **d)** Boxplots representing the PCN per PTU from Gram-negative (Pseudomonadota, purple) and –positive (Bacillota, green) bacteria (Supplementary dataset 2). Only the most abundant PTUs are indicated for each group. Numbers on the right of each boxplot show the number of plasmids belonging to each plasmid group.

Major PTUs and plasmid replication types showed a characteristic PCN (Figure 2d, Supplementary dataset 2, Supplementary Fig. 3). Among the most abundant plasmid groups, HCPs were mainly associated with Col-like replicons in Gram-negatives (e.g., PTUs E9, E10, E3)^22^, and with rolling-circle replicating plasmids in Gram-positives (e.g., PTUs Bac20, Lab37)^23,24^. On the other hand, LCPs were frequently associated with well-characterized Gram-negative enterobacterial plasmids, such as the widespread IncF family (e.g., PTUs F_K_ and F_E_). In Gram-positives, LCPs were diverse and included plasmids related to theta-replicating prototypical plasmids (e.g., PTU-Bac8, PTU-Bac42, PTU-Lab18)^25–28^. Regarding mobility, conjugative plasmids were typically present at low PCNs (median = 2.17), while non-mobilizable plasmids and particularly mobilisable plasmids (median = 3.94 and 8.58, respectively) were associated with a significantly higher PCN (Kruskal-Wallis test followed by Dunn’s test for pairwise multiple comparisons p < 10^-35^, Supplementary Fig. 4).

### PCN is independent of genetic repertoire, bacterial host, and co-resident plasmids

The above results highlight that each plasmid group has a characteristic PCN that is likely a direct consequence of their biology. However, there is also substantial variation in PCN within plasmid groups, at least for some of them (see, for instance, PTU-E9 and PTU-Bac19). To characterise the sources of this variability, we first focused on how gene content and similarity affected PCN. By comparing the PCN of plasmids bearing the same replicon type but belonging to different PTUs, we found that, in general, PCN was conserved in most replicon types regardless of the genetic content (Supplementary Fig. 5, Supplementary Dataset 3).

Analysis of the exceptions revealed interesting plasmid biology features. For instance, Col-like (rep_2335) plasmids from the *Escherichia*-associated PTU-E63 were present at significantly lower PCNs than those belonging to broader host range PTUs (PTU-E3 and PTU-E76; Kruskal-Wallis test followed by Dunn’s test p < 10^-2^). Similarly, IncFIB/IncFII plasmids showed significant, although small, PCN differences between the *Klebsiella*-associated PTU-F_K_ and the *Salmonella*-associated PTU-F_S_ (Kruskal-Wallis test followed by Dunn’s test p < 10^-2^, Supplementary Fig. 5, Supplementary Dataset 3).

Prompted by these observations, we next investigated the impact of host range on PCN. Of the 64 replicon types and 57 PTUs shared between at least two different genera, only four replicons, and 5 PTUs showed significant differences in their PCN across hosts (Supplementary Fig. 6, see Supplementary Dataset 4 for statistical analyses). Similarly, only two of the 13 plasmid clusters present in multiple host genera displayed significant differences in PCN between hosts (Wilcoxon rank sum test, adjusted p < 0.04, Supplementary Fig. 7). Lastly, we investigated how the presence of other plasmids within the cell affects PCN and found that only 10% of the PTUs and 7% of the replicon types showed a statistically significant correlation between PCN and the number of plasmids in the cell (Pearson’s rank correlation, p < 0.049, n = 114 for PTUs and n = 212 for replicon types, Supplementary Dataset 5).

Overall, and in agreement with previous small-scale observations^29,30^, our results suggest that most plasmids encode replication control mechanisms that robustly control PCN independently of the content and identity of the plasmid’s genetic repertoire, the host, and the presence of other co-resident plasmids.

### Intrinsic PCN variability is higher in HCPs

We reasoned that the observed variability in PCN might be a direct manifestation of the stringency of replication control across plasmid lifestyles. In agreement with this hypothesis, HCPs showed significantly greater variability in their PCN than LCPs (measured as coefficient of quartile variation – CQV^31^, Wilcoxon rank-sum test *p* < 10^-9^, effect size ≥ 0.836 and ≥ 0.512 for both replicon types and PTUs, respectively, Figure 3, Supplementary Fig. 8, Supplementary Fig. 9), even after accounting by host shared ancestry (as estimated by Bayesian multilevel models with host as a random effect; see Supplementary Dataset 6 for details). Moreover, intrinsic variability and PCN were positively and strongly correlated when we classified plasmids according to their replicon type, PTU, or plasmid cluster (Spearman’s rank correlation *p* < 10^-2^; Figure 3, Supplementary Fig. 9). As an illustrative example, the PCN of Col-like HCPs varied over one order of magnitude (CQV > 50), while LCPs such as IncF plasmids displayed smaller variations in their PCN (CQV ∼ 15-24; Supplementary Fig. 10).

**Figure 3.**
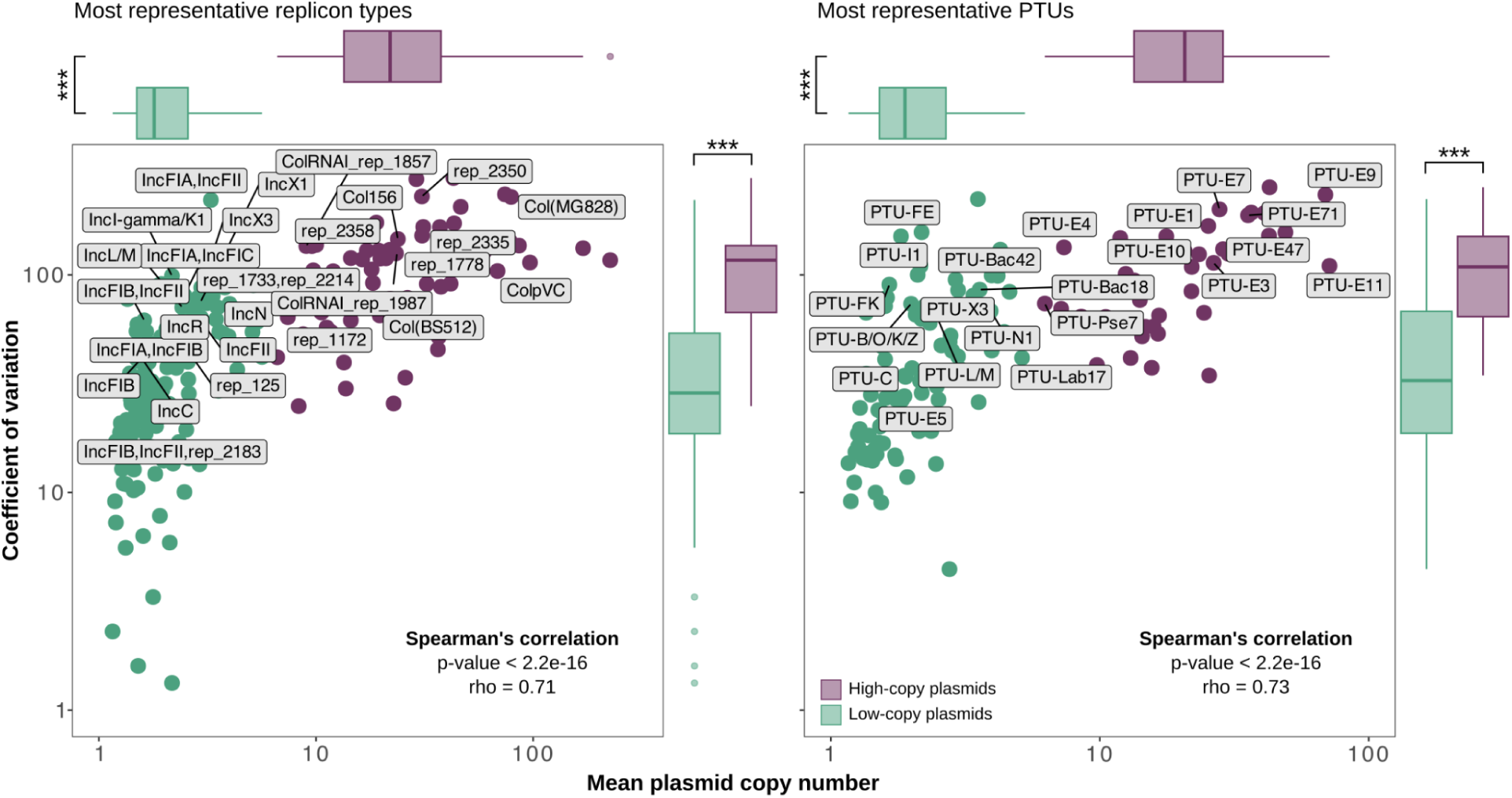
Intrinsic PCN variability. Correlation of PCN (x-axis) per replicon type (left) and PTU (right) with the variability in PCN as measured by the coefficient of quartile variation (CQV; y-axis). Each dot represents the mean PCN and the quartile variation for a plasmid group. The Spearman’s rho and p-value for each correlation are shown at the bottom right corner of each plot, and n indicates the sample size. Labels indicate the most abundant plasmid groups. Colours distinguish HCP (purple) from LCP (green). Boxplots represent aggregated PCN (top) and CQV (right) data for all plasmid groups, according to their classification as HCPs and LCPs. The line inside the box marks the median. The upper and lower hinges correspond to the 25th and 75th percentiles, and whiskers extend to 1.5 times the interquartile range. Asterisks indicate significant differences between HCPs and LCPs (***: p < 10^-9^; Two-sided exact Wilcoxon rank-sum test, effect size ≥ 0.836 for all tests of replicon types and ≥ 0.512 for all tests of PTUs; see Supplementary Dataset 6 for Bayesian multilevel models with host as a random effect).

In contrast to previous observations restricted to model laboratory plasmids^32,33^, these results indicate that the higher the PCN, the more relaxed the control of replication and segregation. As gene expression and PCN are tightly linked^12,13^, this result underscores the role of HCPs as plastic adaptive platforms^15^. On the other hand, biotechnological and synthetic applications may benefit from the reduced noise of LCP-derived vectors to ensure precise control of gene expression.

### Replicon dominance determines plasmid copy number in multi-replicon plasmids

Multi-replicon plasmids are abundant and often occur due to plasmid co-integration, a phenomenon by which two plasmids merge as a single DNA molecule^34^ (Figure 4a). To shed light on whether multiple replicons interact to control PCN, we tested how PCN varies for a given replicon when it drives plasmid replication alone (single replicon form) or when it co-exists with other replicons within the same plasmid molecule (multi-replicon form). We found 51 replicon types present in both forms. Of those, 37% (19/51) showed significantly different PCN between the single and multi-replicon forms (Wilcoxon rank sum exact test p < 0.047 in all cases; see Supplementary Fig. 11 for data represented as boxplots). Some replicons (e.g., IncQ1, Col156, or IncFII) exhibited a lower PCN when present in multi-replicon plasmids, while other replicons showed higher PCN (e.g., ColE1-like replicons rep_2358 and rep_2370; Figure 4b). This demonstrates that interactions between co-existing replicons frequently alter PCN.

**Figure 4.**
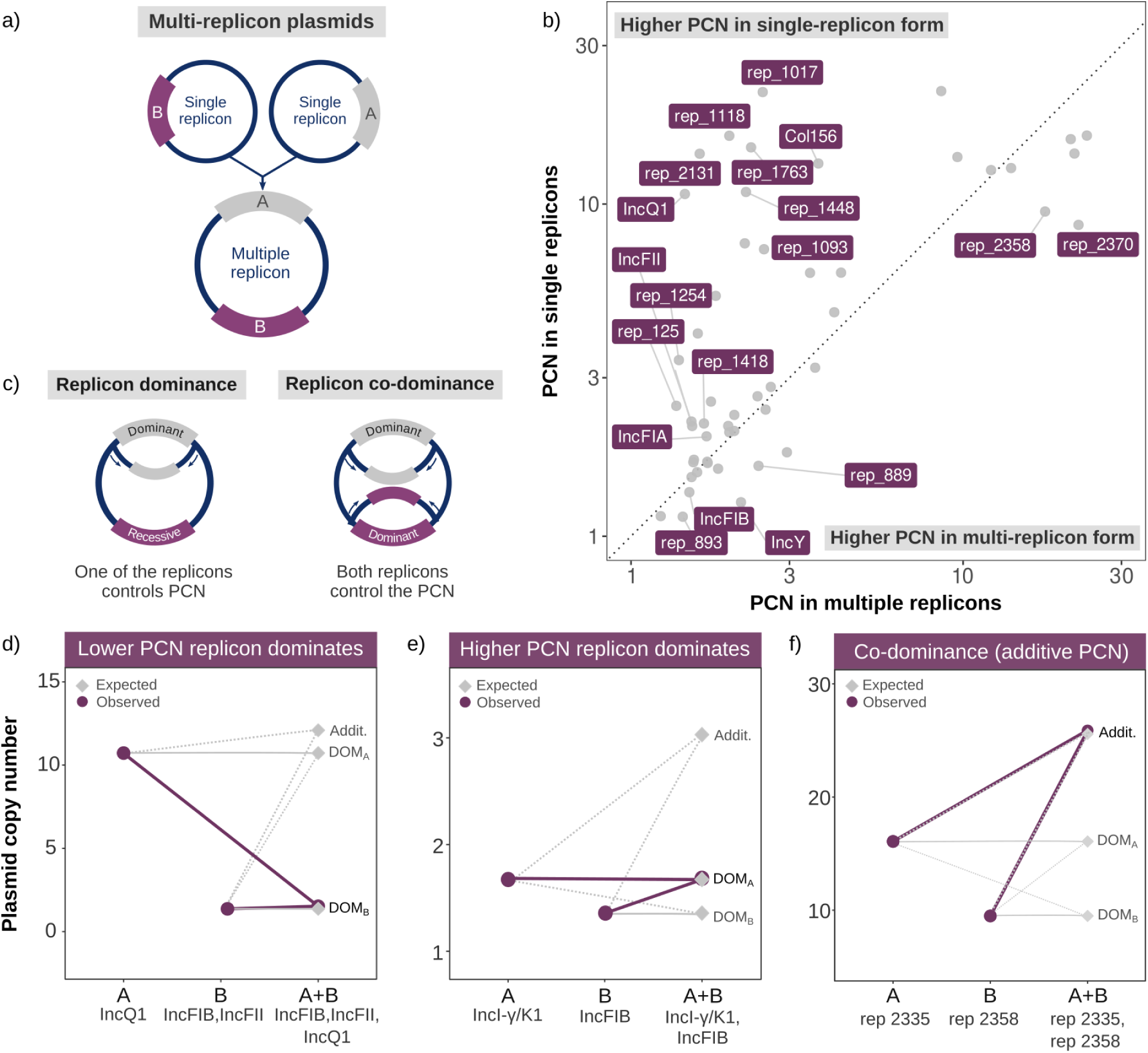
Replicon dominance determines the plasmid copy number of multi-replicon plasmids. **a)** Scheme depicting the formation of a multi-replicon (co-integrated) plasmid. **b)** Acquisition of additional replicons frequently alters PCN. For each replicon type, the PCN of plasmids as single-replicon (y-axis) is plotted against their PCN when co-existing with other replicons in a multi-replicon plasmid (x-axis). See Supplementary Fig. 11 for the same data represented as boxplots. The dotted line indicates no change in PCN between forms. Replicon types with significant PCN differences between single– and multi-replicon forms are labelled (p < 0.047; Two-sided exact Wilcoxon rank-sum test with continuity correction). **c)** Mechanisms controlling PCN in multi-replicon plasmids. The diagram on the left depicts *replicon dominance*, where one replicon (dominant) exerts full control over the PCN, while the other (recessive) has no influence. The diagram on the right depicts *co-dominance* between the two replicons, where both contribute to the final PCN. **d)** Example of a multi-replicon plasmid where the replicon with lower PCN (IncFIB, IncFII; replicon B) is dominant over the replicon with higher PCN (IncQ; Replicon A). **e)** Example where the higher PCN replicon (IncI-γ/K1; replicon A) is dominant over the one with lower PCN (IncFIB; replicon B). **f)** Example of co-dominant replicons. Grey diamonds on panels **d-f** show predicted dominance outcomes: PCN additivity (Addit.), dominance of the higher (DOM_A_) or lower (DOM_B_) PCN replicon. Observed data (purple circles) depicts the observed median PCN for each case. Raw data for each replicon is available in Supplementary Fig. 12.

To explain these interactions, we borrowed from classical genetics and conceived the concept of *replicon dominance*. We observed that in certain replicon combinations, one of the replicons did not contribute to the final number of plasmid copies (i.e., it was recessive), and the PCN was controlled by the other replicon(s) (i.e., dominant) (Figure 4c). Higher copy number replicons were generally recessive to replicons showing lower copy numbers (6 of 22 cases; Figure 4d, Supplementary Fig. 12). For instance, HCP-associated replicons were recessive to LCP replicons, possibly because their replication mechanism (e.g., strand displacement) is unsuitable for efficiently replicating larger plasmids (Figure 4d, Supplementary Fig. 12, Supplementary Fig. 13). On the other hand, the higher copy replicon only dominated in plasmids containing multiple LCP-associated replicons, albeit PCN differences were generally small (2 of 22 cases; Figure 4e, Supplementary Fig. 12, Supplementary Fig. 13).

An interesting case of replicon dominance occurs when both replicons are co-dominant, resulting in an additive PCN (2 of 22 cases, Figure 4c,f, Supplementary Fig. 12, Supplementary Fig. 13). We found co-dominance exclusively in plasmids carrying Col-like replicons, indicating that it might be a specific feature of plasmids of this group. Indeed, a relatively small number of mutations can lead to additive PCN in single-replicon co-existing Col-like plasmids^30,35^, suggesting that independence (orthogonality) between plasmid replication systems explains replicon co-dominance (Supplementary Figs. 12-14). We also observed other interactions, such as incomplete dominance (4 of 22 cases), resulting in intermediate PCN, antagonism between replicons (1 of 22 cases), and high-order interactions occurring in plasmids showing more than two replicons (7 of 22 cases). However, due to weak statistical support or the small number of cases, we refrain from discussing them in detail (Supplementary Fig. 12).

### A pervasive scaling law rules plasmid biology

Next, we sought to identify which factors determine PCN using a random forest regression model. Random forest regressors are supervised machine learning algorithms that leverage ensembles of decision trees to predict continuous variables. To train and refine our model, we used numerical and categorical variables from our dataset (see methods). The model could predict PCN using these variables, although with modest performance (Supplementary Fig. 15, Mean Absolute Error (MAE): 5.18, R^2^: 0.51). Interestingly, plasmid size was the variable that held more predictive power in our dataset (Gini feature importance = 40%), well above other features typically associated with PCN (e.g., PTU or plasmid mobility; Gini feature importance ≤ 10%; Supplementary Fig. 15).

Indeed, although there was substantial unexplained variance, plasmid size and copy number were strongly and negatively associated (Supplementary Fig. 16), and their relationship followed a power law (being linear in a log-log plot; Figure 5a). Power laws are typically defined by the formula *y = a · x^b^*, which, in this case, takes the following form: *PCN = 10^c^ · size^k^*, where *c* is the intercept, and *k* is the scaling factor or slope. The overall slope was *k =* –0.65 (95% CI: –0.66, –0.63), indicating that, on average, a 1% increase in size is associated with a 0.65% decrease in PCN (Supplementary dataset 7).

**Figure 5.**
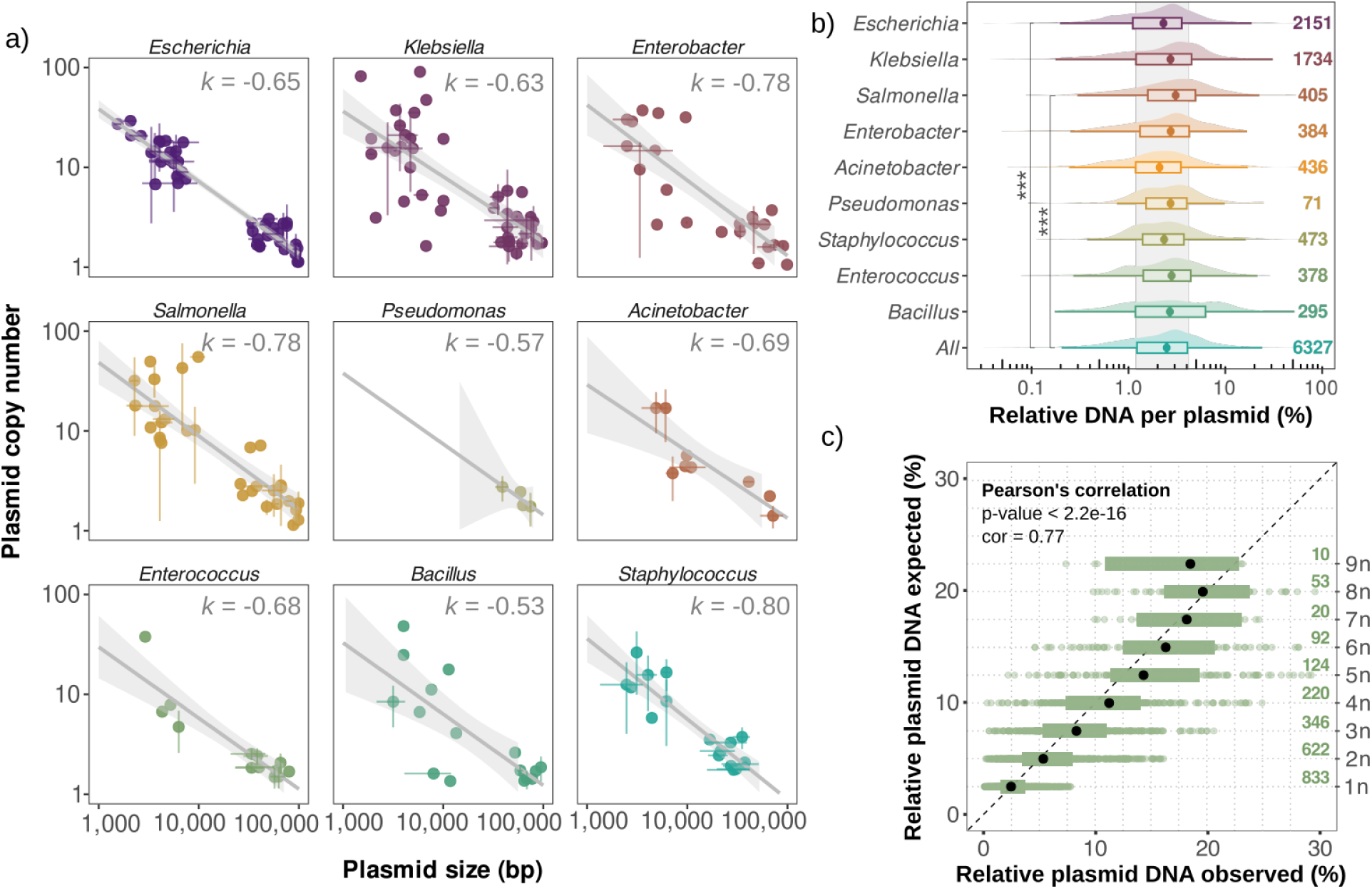
A scaling law links copy number and plasmid size across bacterial phylogeny. **a)** Scatter plots showing the correlation between plasmid size (x-axis) and PCN (y-axis) for the analyzed genera. Each point represents the median PCN and plasmid size for each PTU, and error bars indicate the standard deviation from the median. Grey lines represent ordinary least squares regression, with the surrounding shaded area indicating 95% confidence intervals. The scaling factor or slope, *k*, is indicated on each panel. **b)** Distribution of total DNA load per plasmid (x-axis) relative to chromosome size per genus (y-axis). The DNA load of each plasmid is calculated by multiplying the plasmid size by the copy number and then expressed as a proportion relative to the chromosome size. The point inside the box marks the median. The upper and lower hinges correspond to the 25th and 75th percentiles, and whiskers extend to 1.5 times the interquartile range. Only *Escherichia* and *Salmonella* significantly differ from All; Kruskal-Wallis test followed by Dunn’s test for pairwise multiple comparisons p < 10^-4^; effect size = 0.006. **c)** Relative plasmid DNA load observed (%) (x-axis) and expected (y-axis) per cell. The y-axis indicates the expected plasmid DNA load (%) inside a cell when it contains one plasmid (1n), two plasmids (2n), and so on. This expected data has been calculated by generating a sequence from 1 to 9 multiplied by the median of the DNA load per plasmid (2.49%). Each green point represents a single genome, and the black points are the median for each category. Shading indicates interquartile ranges. Pearson’s p-value and coefficient are shown for the correlation between expected and observed plasmid DNA.

This suggests that a scaling law drives the relationship between size and PCN. Scaling laws are prevalent in many natural systems, revealing patterns and relationships across orders of magnitude. In biology, scaling laws typically take *k* values of ⅔ or ¾ and can be leveraged as powerful tools for modelling and understanding complex systems^36,37^. Some examples of scaling laws include the relationship between metabolic rate and body size in animals and plants^38,39^ and the scaling of gene content with regulatory networks in bacterial genomes^40^.

To test the universality of the *k* ≈ –0.65 (i.e., *k* = –⅔) relationship, we calculated *k* for each genus in our dataset. Although plasmids from Gram-negative and –positive bacteria (in this case belonging to the Pseudomonata and Bacillota phyla) are very diverse, copy numbers and plasmid sizes scaled similarly. Indeed, all genera presented slopes not significantly different to *k* = *-*0.65 (One-sample t-test, BH adjusted p > 0.51 in all cases, Figure 5a, Supplementary Fig. 17 and Supplementary dataset 7). The conservation of *k* values across bacterial groups further highlights the universality of the PCN-size scaling law and provides a simple formula to roughly estimate the PCN of any plasmid (see methods).

### Plasmid DNA load is conserved relative to chromosomal size

To shed light on the metabolic constraints imposed by plasmids, we calculated the total DNA content of each plasmid as the product of copy number and size. This reflects the total amount of DNA (in bp) of a given plasmid within a cell or its *DNA load*. Plasmids from *Pseudomonas*, *Bacillus, Salmonella*, and *Klebsiella* accounted for greater DNA loads than average, while the reverse was observed for plasmids from *Acinetobacter*, *Staphylococcus*, and *Enterococcus* (Supplementary Fig. 18, Kruskal-Wallis test followed by Dunn’s test for pairwise multiple comparisons p < 10^-3^).

We wondered whether variation in chromosome size could explain differences in plasmid DNA load across genera, particularly given that chromosome and plasmid size correlate^2,41^. Although there was substantial variation, our analyses revealed that, regardless of their host genus, size, or copy number, all plasmids tended to account for approximately the same percentage of chromosomal DNA (median = 2.49 %, IQR: 1.22-4.06, Figure 5b, only *Escherichia* and *Salmonella* being significantly different, albeit negligibly from All; Kruskal-Wallis test followed by Dunn’s test for pairwise multiple comparisons p < 10^-4^; effect size = 0.006). This conserved *relative plasmid DNA load* indicates that common constraints control the interplay between copy number and size in HCPs and LCPs.

Given that PCN is independent of co-resident plasmids (Supplementary Dataset 7), we checked whether relative plasmid DNA load scales proportionally to the number of plasmids within the cell. Thus, if any given plasmid accounts for ∼2.5% of the genome, the cumulative plasmid DNA load in a cell would be the product of that DNA fraction by the number of plasmids. As such, a cell harbouring two different plasmids would have a relative plasmid DNA content of 2n (∼5%), a cell with three plasmids would have 3n (∼7.5%), and so on. Remarkably, this expectation correlates well with the observed percentage of DNA content allocated to plasmids (Pearson product-moment correlation r = 0.77, p < 10^-6^; Figure 5c). Therefore, the proportion of plasmid DNA within any bacterial cell, indeed, seems to follow a discrete pattern.

## Discussion

Copy number is an essential feature of plasmid biology. PCN not only determines a fundamental division between plasmid lifestyles but also drives key differences in gene expression, metabolic burden, and antibiotic resistance^1,14,15,42^. In this work, we leveraged sequencing data to obtain, for the first time, a large-scale dataset of the copy number of 6,327 plasmids (Figure 1). We found that PCN varied widely, ranging from ∼1 to more than 1,000 copies per cell and that it was generally bimodally distributed. This reflects two well-known plasmid lifestyles, for which a clear distinction was lacking^18,43,44^ (Figure 2). PCN was generally independent of the content and identity of the plasmid’s genetic repertoire (Supplementary Fig. 5), the presence of co-resident plasmids (Supplementary Dataset 5), and the bacterial host (Supplementary Fig. 6 and 7). In line with previous observations^4^, these results emphasise that intrinsic replication control mechanisms are crucial in determining each characteristic PCN, but also provide new insights into how these mechanisms differ among plasmid families. The stringency of PCN control is, however, different between plasmid lifestyles: for instance, HCPs show more variation in PCN than LCPs (Figure 3). In addition, we devised the concept of replicon dominance and used it to explain the interactions defining PCN in widespread multi-replicon plasmids (Figure 4). In this regard, perhaps the most relevant result is that low PCN replicons are generally dominant to high PCN replicons. This suggests two non-mutually exclusive possibilities that await experimental validation: i) the replication machinery of the HCP (small) plasmid is inefficient for replicating a larger DNA molecule, and/or ii) selection favours larger plasmids that exist in low copy, as a mechanism to reduce fitness costs.

Arguably, the most intriguing result of our work is that a PCN-size scaling law governs plasmid biology across bacterial species (Figure 5). This result agrees with previous observations with limited sampling^10,11^ or concerning only Enterobacterial plasmids^9^ and is further supported by a recent work identifying consistent scaling laws that relate plasmid size with copy number, protein-coding genes, and metabolic genes across ecological niches^45^. Altogether, these complementary works indicate that universal constraints orchestrate the PCN-size trade-off. However, the underlying molecular mechanism remains to be uncovered. Plasmid replication might be constrained by a limitation in cellular resources, such as metabolites (e.g., nucleotides), cell machinery (e.g., polymerases and helicases), or even physical intracellular space. Nevertheless, we found that the presence of multiple co-resident plasmids does not affect PCN and that each plasmid independently accounts for a similar DNA load (∼2.5% of the chromosome size; Figure 5). This suggests that rather than the availability of cellular resources, the efficiency (e.g., replication rates or the turnover of assembled replisomes)^46^, regulation (e.g., in response to cellular biomass, cell cycle, or culture growth phase)^47^ or timing (e.g., synchronicity with the cell-cycle)^48,49^ of biophysical processes within the cell might explain the PCN-size scaling law.

Our study is not without limitations. First, PCN estimation might be subject to a certain degree of noise. PCN is an inherently plastic trait and may vary at different points of the host cell cycle or depending on growth conditions^10^. We calculated PCN from deposited sequencing data and cannot exclude that some experimental factors (e.g., sequencing technology, DNA extraction protocol) may affect PCN determination^50–52^. Second, some plasmids in our database may be synthetic, and consequently, they might have been engineered to display an artificially high (or low) PCN. Third, our results derive only from a few bacterial taxa, primarily genera of clinical importance. To some extent, this is an unavoidable consequence of the lack of appropriate tools for plasmid classification (e.g., replicon type, PTU) beyond well-studied bacterial genera. Fourth, our analyses rely on the accuracy of these bioinformatic tools for establishing meaningful plasmid classifications. While these methods are standard in the field, they could inadvertently introduce bias by mispredicting some plasmid properties (e.g., mobility)^53^. Although these factors probably account for some of the observed variability in PCN, they are unlikely to significantly impact our general conclusions, founded upon an analysis of thousands of diverse bacterial plasmids.

Finally, our analysis is restricted to the classical definition of plasmids (independently replicating circular DNA molecules). Yet, not all plasmids are circular, and many extrachromosomal genetic elements share properties with plasmids (e.g., phage-plasmids or secondary chromosomes)^54^. In this sense, our study lays the foundation for future works addressing copy number variation in extrachromosomal genetic elements of non-model microorganisms. By revealing a traditionally neglected aspect of their biology, these studies will shed light on the complex interplay among different genetic elements and their bacterial hosts.

In summary, our comprehensive analysis uncovers the principles that drive PCN. From an applied perspective, leveraging these principles will enhance the design of plasmids as biotechnological tools (e.g., noise in gene expression, stability of large constructs, optimization based on host chromosome size). Further, we provide a method to predict PCN, which will be useful to, for instance, improve the assembly of plasmid sequences from metagenomic samples. From a fundamental standpoint, our study provides a detailed catalogue of PCNs across plasmid groups, highlighting the major sources of variability and paving the way for understanding the fundamental constraints that govern plasmid biology.

## Methods

### Data processing

To build our database of complete, high-quality plasmids and their PCN, we focused on nine different genera from the phyla Pseudomonadota and Bacillota (i.e., Gram-negative and –positive bacteria). Specifically, we selected the following genera: *Acinetobacter*, *Bacillus*, *Enterobacter*, *Enterococcus*, *Escherichia*, *Klebsiella*, *Pseudomonas*, *Salmonella* and *Staphylococcus*, which include species with biotechnological and clinical interest, such as all members of the ESKAPEE group^19^. We identified and downloaded all available assemblies from the selected genera annotated as Complete Genomes in the NCBI database (n = 24,674) on 5/12/2023. SRA information was extracted using the sra-toolkit v2.11.3 (https://github.com/ncbi/sra-tools) with a custom pipeline (https://github.com/PaulaRamiro/NpAUREO/) and used to download available paired-end reads (corresponding to n = 3,156 assemblies).

Reads were aligned against their respective assemblies to extract the trimmed-mean coverage using CoverM v0.6.1 (https://github.com/wwood/CoverM) with the following command: *coverm contig –m trimmed_mean.* Some of the alignments did not meet the quality criteria of CoverM and were excluded from further analyses (n = 678). Plasmids were identified using mob_suite (see ***Plasmid classification*** for details; n = 8,660 plasmids belonging to 2,478 assemblies), and their topology (circular or linear) was checked with a custom script that retrieves information from the NCBI database using its dedicated API (see Code Availability section). Plasmid contigs annotated as circular were kept for further analyses (n = 8,091). The PCN was then calculated for each sample as the ratio between the mean coverage of plasmid contigs and the mean coverage of the chromosome. We removed plasmids belonging to assemblies with an absolute sequencing depth below 30x (n = 736) and plasmids showing a size < 1 kb (n = 28) or PCN <1 (n = 1,000). As a quality control, we confirmed that the PCN values calculated using CoverM were consistent with those reported in other studies, showing a strong correlation between different methods (Supplementary Fig. 19)^9,55,56^. This approach led to a final dataset of 6,327 plasmids and their PCN (Supplementary Dataset 1).

### Plasmid classification

We classified plasmids using several complementary methods. First, we typed plasmids into different incompatibility groups according to their replication mechanism^5,20^ using MOB-typer from mob_suite v3.1.8 (https://github.com/phac-nml/mob-suite)^20^ using the flag *--multi* to type independent plasmids within samples. This method leverages features in the DNA sequences responsible for plasmid replication (e.g., encoding replication initiation proteins) to establish plasmid groups whose replication is mechanistically similar, termed replicon types. Second, we used a classification scheme based on similarity across the whole plasmid genetic content with COPLA v1.0^21^. Plasmids that share high homology (>70%) in more than 50% of their sequence are assigned to the same plasmid taxonomic unit (PTU)^21,57^. Although PTUs and replicon types were strongly associated (Supplementary Fig. 20), we could assign a replicon type to nearly 90% of the plasmids, but only 63% belonged to defined PTUs (Supplementary Dataset 2). Indeed, nearly 4% of the plasmids belonged to new, still unnamed PTUs, while the rest (32%) could not be accurately classified.

To further complement these classifications, we employed a custom clustering approach: Plasmid sequences were extracted from the FASTA files of the assemblies and annotated with Bakta v1.9.3 (https://github.com/oschwengers/bakta)^58^. A distance matrix using gene-by-gene presence-absence was created using the *accnet* function of PATO v1.0.6 (https://github.com/irycisBioinfo/PATO)^59^ with a Jaccard distance similarity parameter of 70%. Then, we generated a k-nearest neighbours network (K-NNN) to allow reciprocal connections with k = 10 neighbours. Plasmids were clustered from the K-NNN using mclust^60^ v6.1.1. Finally, we also used MOB-typer (with the *--multi* flag) to predict plasmid mobility. We note, however, that this method likely overestimates the fraction of plasmids assigned to the non-mobilizable category^53,61^.

### PCN Analysis

All analyses were performed in R (v4.1.2). Analysis of the modes for PCN distributions was conducted by first checking the number of modes of the distribution with LaplacesDemon^62^ v16.1.6 R package and then using the *locmodes* function from the R package multimode^63^ v1.5, which estimates the locations of both modes and antimodes, with default parameters. To measure PCN variation across our dataset, we calculated the quartile coefficients of dispersion (CQV). The CQV allows for robustly comparing the degree of variation from one plasmid group to another, even if the PCNs are drastically different^31^. CQV was calculated with the R package cvcqv^64^ v1.0.1. Plasmid DNA load (bp) was calculated as *plasmid load (bp)* = *plasmid size (bp)* × *PCN*. The relative percentage of plasmid DNA load was calculated as 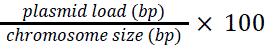. Taken together, the relative percentage of plasmid DNA can also be expressed as 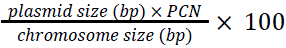.

### Replicon dominance analysis

To visualize the statistical significance of differences in PCN between groups, we employed a letter-based classification using the *cldList* function from the rcompanion^65^ v2.4.36 R package. *cldList* was used to assign letters to each group based on the p-values for the Dunn test performed after the Kruskal-Wallis test (see Statistical analysis and regression). Groups without statistically significant differences were excluded from the analysis. In cases where the multi-replicon form was not different from only one of the simple replicons (e.g. ‘a’, ‘a’, ‘b’), it was identified as a case of *dominance*. Other occurrences, such as (’a’, ‘ab’, and ‘b’), or ( ‘a’, ‘b’, and ‘c’), were classified as other interactions.

We then checked cases where the multi-replicon form had a higher median PCN than each single form to find cases of *co-dominance*. In those cases, to obtain statistical support and test for an additive effect, we generated a bootstrapped distribution representing the sum of the single replicons and compared it to the observed values. We excluded *co-dominance* cases when the PCN of the multi-replicon and that of the bootstrap were significantly different.

### Model training and formula usage

To train the prospective model, manual curation of the dataset was performed first to remove redundant or non-informative variables for PCN (e.g., species). Also, categorical variables with too many classes were eliminated if other variables contained the same information with fewer classes. The final list of variables used to train the model was as follows: genus, predicted mobility, the presence of single or multiple replicons, size of the chromosome of the host, GC content of the plasmid, number of plasmids present in the host, predicted PTU, and size of the plasmid (with a log_10_ transformation). The output variable was PCN. Observations with PCNs > 100 were considered outliers and eliminated. Observations with unknown or unassigned PTUs were also eliminated to suppress noise in the dataset.

Several models, including scikit-learn v1.5.2 simple linear regression^66^, generalised linear models^66^, elastic net regressor^66^, multi-layer perceptron regressor^66^, random forest regressor^66^ and XGboost^67^ v2.1.1, were pre-tested with light tuning. In all trials, sklearn-RandomForestRegressor outperformed all other models. After selecting RandomForestRegressor, further tuning was performed: first, a random search cross-validation with a wide parameter range (initial parameters available in the code repository), and then, a deeper grid search cross-validation with values around the parameters selected in the random search. Finally, recursive feature elimination was used to improve the final model using mean absolute error as the performance metric. Gini Feature importance was directly extracted from the model using the built-in function. The complete code and dataset used for the final model are available at https://github.com/PaulaRamiro/NpAUREO/tree/main/Model.

### Statistical analyses

The significance level was set at 0.05 for all statistical tests. All statistical tests performed were two-tailed. In all boxplots, the box size extends to the interquartile range (IQR), and the line represents the median. Whiskers extend from the edges of the box to the smallest and largest values within 1.5 times the IQR. Outliers are plotted as individual points beyond the whiskers. The Chi-squared test was used to compare the counts of HCPs and LCPs of each genus against the total population to reveal significant over or underrepresentation of either. Cohen’s h was calculated using the corresponding formula: 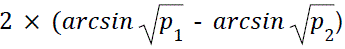, where *p*_1_ and *p*_2_ are the proportions being compared.

When data did not meet the assumptions for a one-way ANOVA (normal distribution and homoscedasticity), the Kruskal-Wallis test was used to compare multiple groups. Effect size was calculated as Eta squared using kruskal_effsize from rstatix v0.7.2 R package^68^. Dunn’s test was further performed to determine which groups presented statistically significant differences.

To compare two single independent groups, we employed the Wilcoxon rank-sum test with continuity correction. In cases of multiple testing, we used the Benjamini-Hochberg (BH) correction to control for the false discovery rate. To measure effect size in Wilcoxon rank-sum tests, we employed wilcox_effsize from rstatix^68^.

Bayesian multilevel models were conducted using the brms^69^ package (v2.22.0) in R to examine the impact of plasmid classification on both the coefficient of quartile variation (CQV) and the mean plasmid copy number. In each model, plasmids classification (HCP vs. LCP) was included as a fixed effect, and random intercepts were incorporated for the presence of each genus into the group (*Escherichia, Klebsiella, Enterobacter, Bacillus, Enterococcus, Pseudomonas, Salmonella, Staphylococcus* and *Acinetobacter*) to account for genus-specific variability. Model fitting utilised default priors: fixed effects were assigned weakly informative normal priors (N(0,10)) while group-level effects were given default priors for variance components (commonly a half-Student’s *t* distribution) that constrain these parameters to be positive; the residual standard deviation was also estimated under a default weakly informative prior. Markov Chain Monte Carlo (MCMC) sampling was performed using Stan’s No-U-Turn Sampler (NUTS) with 4 chains run for 2000 iterations each, including 1000 iterations for warm-up (burn-in), yielding a total of 4000 post-warm-up draws. Convergence was assessed through trace plots, Rhat values (which were approximately 1.00 for all parameters), and effective sample sizes. Parameter estimates were summarized with posterior means, 95% credible intervals, probabilities of direction (pd), and the percentage of the posterior distribution within the region of practical equivalence (ROPE) using the *summary* and *estimate_contrasts* functions.

Spearman correlation analysis was used in all cases where the assumptions of Pearson correlation (continuity, linearity, heteroscedasticity, and normality) were unmet. Otherwise, Pearson’s correlation was used. For regressions regarding the scaling law, given the assumption that the source of error is predominantly the dependent variable (PCN) rather than the independent variable (plasmid size), we employed ordinary least squares (OLS) regression for all fits of log-transformed data. This approach is consistent with other published analyses for this type of data^70,71^. The median PCN per each PTU was used to calculate the slopes for each genus. To assess the statistical significance between the slopes of different genera, we fitted OLS with an interaction term for Genus and performed pairwise comparisons among all genera. The formula PCN = 10*^c^* · size*^k^* allows the estimation of PCNs by simply substituting *c* and *k* for the values provided in Supplementary Dataset 7. If the host of the plasmid is unknown, the general values (*c* = 3.4759, *k* = –0.6466) are to be used. However, if the host of the plasmid is known, more precise values are provided to slightly improve the predictions of some genera. The performance of all formulas for our dataset is provided in Supplementary Dataset 7.

## Data availability

The data generated and/or analyzed during the current study are provided in the Supplementary Information and have been deposited in the Zenodo database and can be downloaded from the following repository^72^: https://zenodo.org/records/14979970 and in the GitHub repository (https://github.com/PaulaRamiro/NpAUREO/).

## Code availability

The source code used to run the analyses and produce the results presented in this manuscript is available from ref.^72^, at https://github.com/PaulaRamiro/NpAUREO/ or https://zenodo.org/records/14979970.

## Acknowledgments

We thank Teresa M. Coque, Hildegard Uecker, and Francisco Dionisio for their suggestions. Work in the evodynamics lab (https://evodynamicslab.com/) is supported by project no. PI21/01363, funded by the Carlos III Health Institute (ISCIII) and co-funded by the European Union; CIBER –Consorcio Centro de Investigación Biomédica en Red-(CB21/13/00084), Instituto de Salud Carlos III, Ministerio de Ciencia e Innovación and Unión Europea – NextGenerationEU; Convocatoria SEIMC-FUNDACIÓN SORIA MELGUIZO de Investigación 2021; and funded by the European Union (ERC, HorizonGT, 101077809). Views and opinions expressed are, however, those of the author(s) only and do not necessarily reflect those of the European Union or the European Research Council Executive Agency. Neither the European Union nor the granting authority can be held responsible for them. PR-M is a recipient of a predoctoral PFIS grant (grant no. FI22/00265) from the Carlos III Health Institute (ISCIII), through the Recovery, Transformation and Resilience Plan and Next Generation EU from the European Union. JR-B acknowledges support by a Miguel Servet contract from the Carlos III Health Institute (ISCIII) (grant no. CP20/00154), co-founded by the European Social Fund, ‘Investing in your future’. VFL acknowledges support by a Miguel Servet contract from the Carlos III Health Institute (ISCIII) (grant no. CP22/00164), co-founded by the European Social Fund, ‘Investing in your future’.

## Author contributions

PR-M and IdQ analyzed the data and created the figures. VFL, JAG, and JR-B provided technical support and conceptual advice. PR-M and JR-B conceived the project. JR-B supervised the project. All authors discussed and provided critical feedback during the analysis of the results. All authors wrote, edited, and reviewed the manuscript.

## Supplementary Information for

**Supplementary Figure 1.**
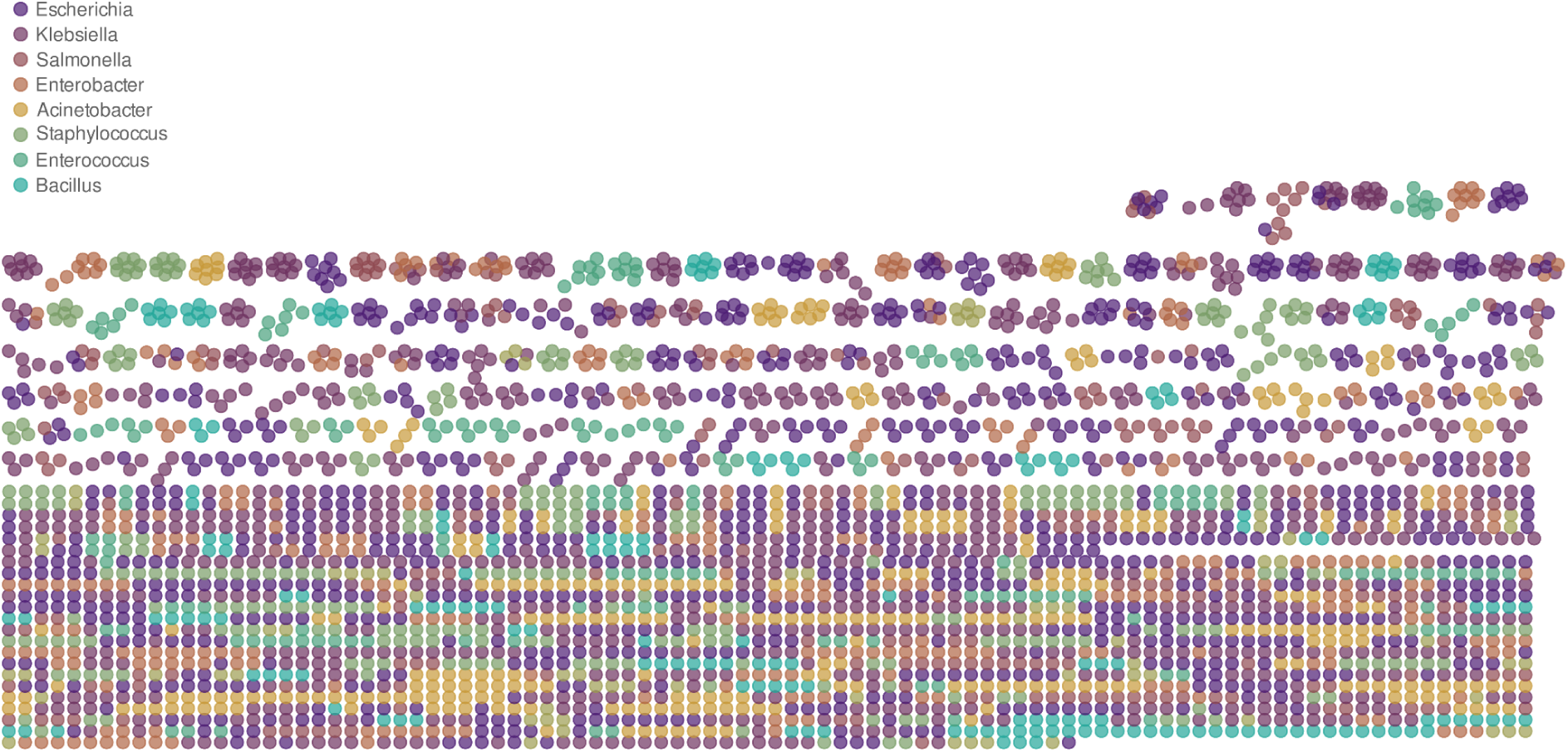
Plasmid clusters containing less than ten members. Plasmid cluster network built with the k-nearest neighbours’ algorithm (K-NNN). Each node represents a plasmid, coloured according to the genus of the bacterial host. Only clusters with less than ten plasmids are shown.

**Supplementary Figure 2.**
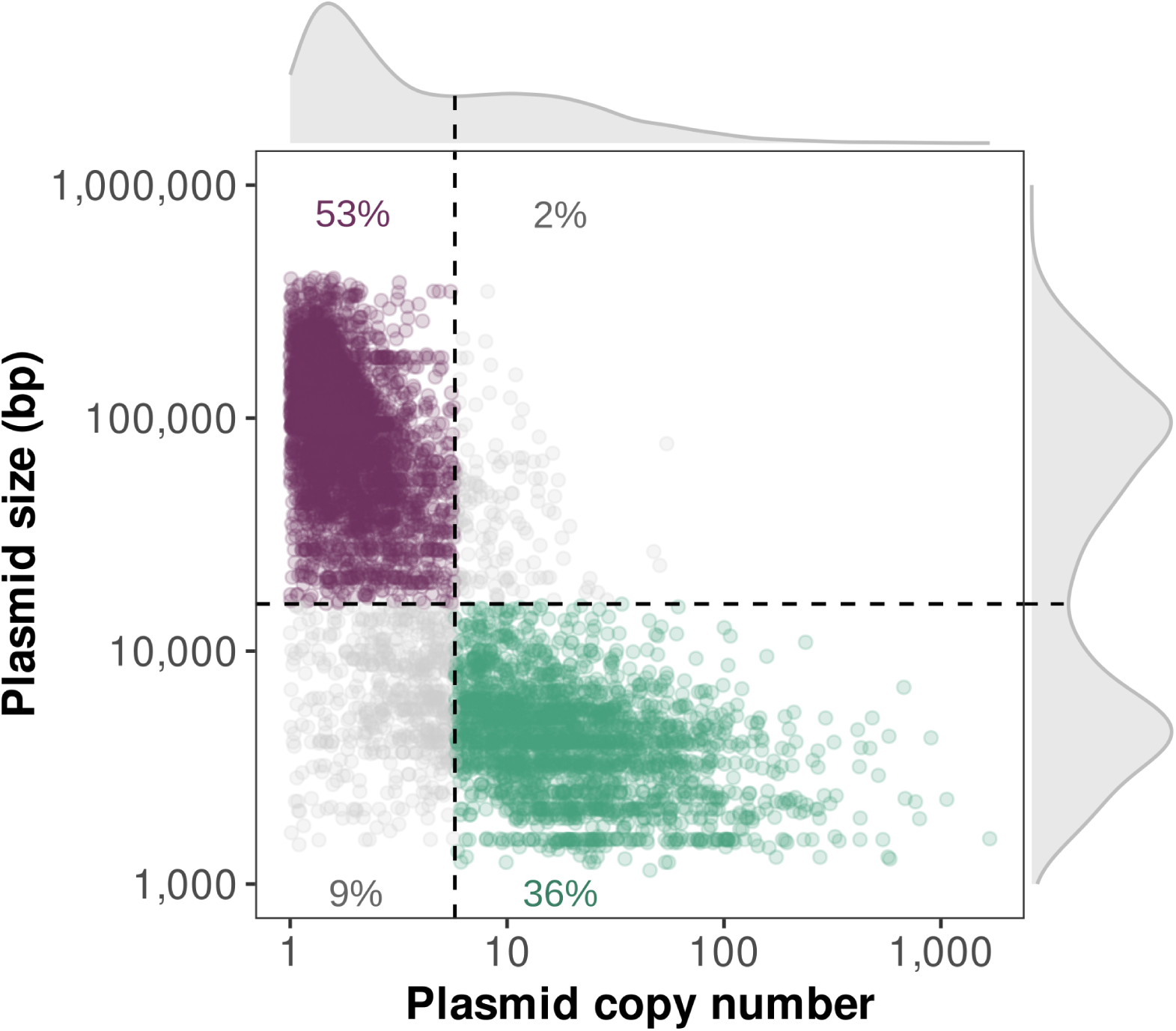
Distribution of plasmids according to size and copy number. Plasmid size (y-axis) and plasmid copy number (x-axis) distributions are shown outside the box. The anti-modes of the distributions for the logarithm (in base 10) of plasmid size (15,939 bp) and plasmid copy number (5.75 copies) are depicted with dashed lines. These limits classify each plasmid in the dataset (individual points) according to its size and copy number. Large LCPs (purple) are represented in the upper-left quadrant, while small HCPs (green) are in the bottom-right quadrant. Large HCPs and small LCPs are depicted in gray in the upper-right and bottom-left quadrants, respectively.

**Supplementary Figure 3.**
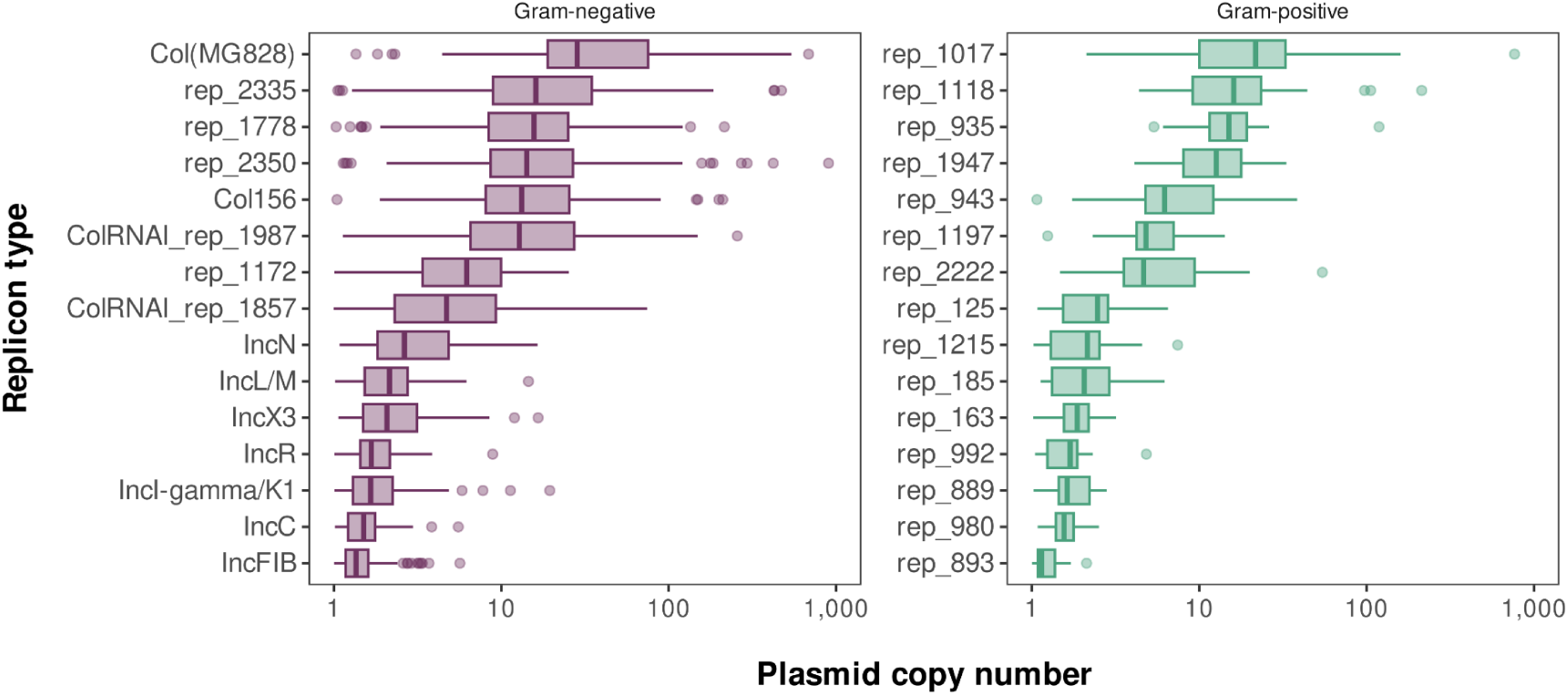
Plasmid copy number variability within replicon types. The boxplots represent plasmid copy number (x-axis) per replicon type (y-axis) in Gram-negative (left panel) and Gram-positive (right panel) bacteria.

**Supplementary Figure 4.**
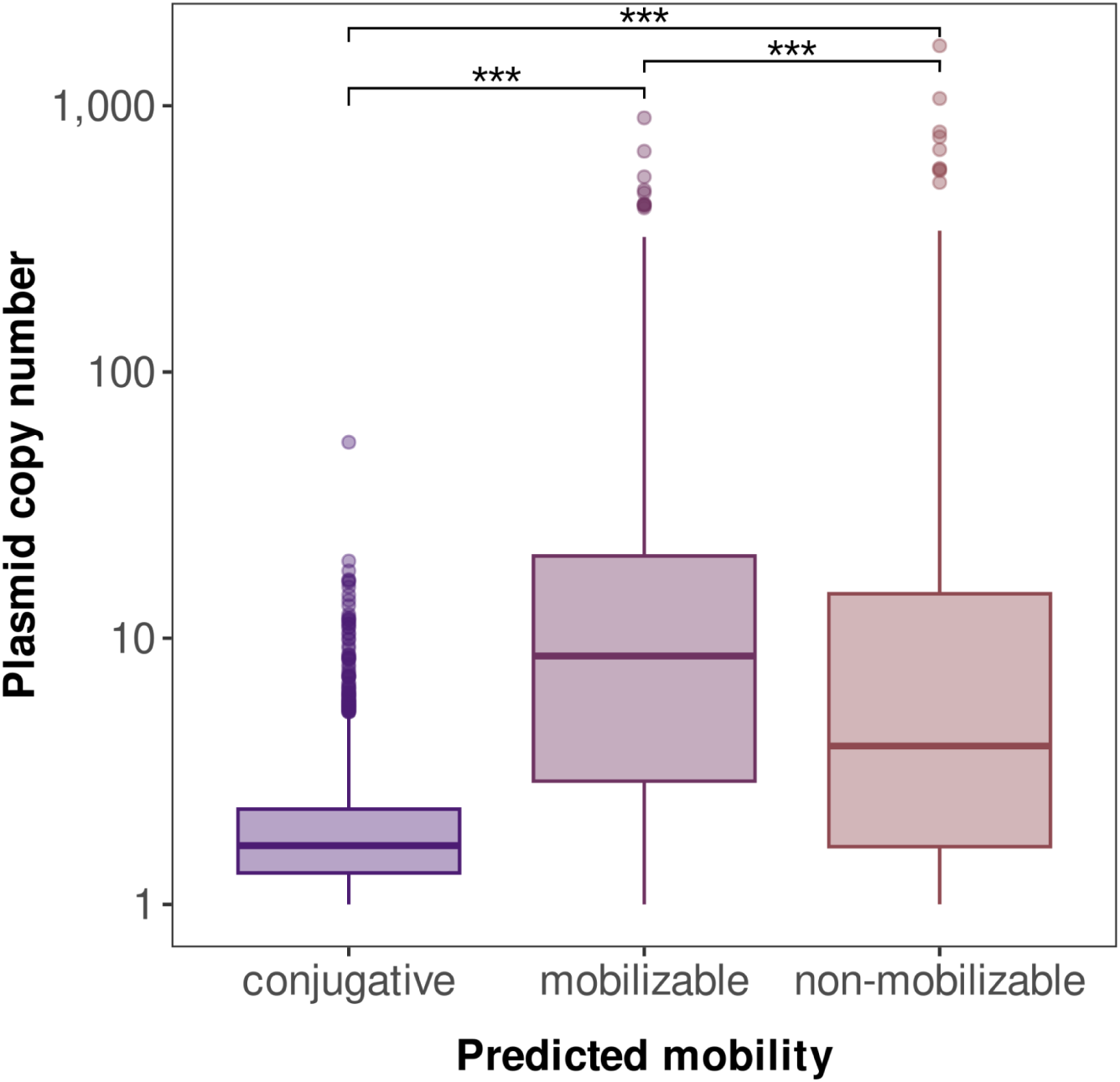
Copy number according to plasmid’s predicted mobility. The y-axis represents the plasmid copy number, while the x-axis represents plasmid categories based on predicted mobility: conjugative, mobilizable, and non-mobilizable. Each boxplot shows plasmid copy numbers within each category. Statistical significance between groups is indicated by asterisks (Kruskal-Wallis test followed by Dunn’s test for pairwise multiple comparisons, p < 10^-5^).

**Supplementary Figure 5.**
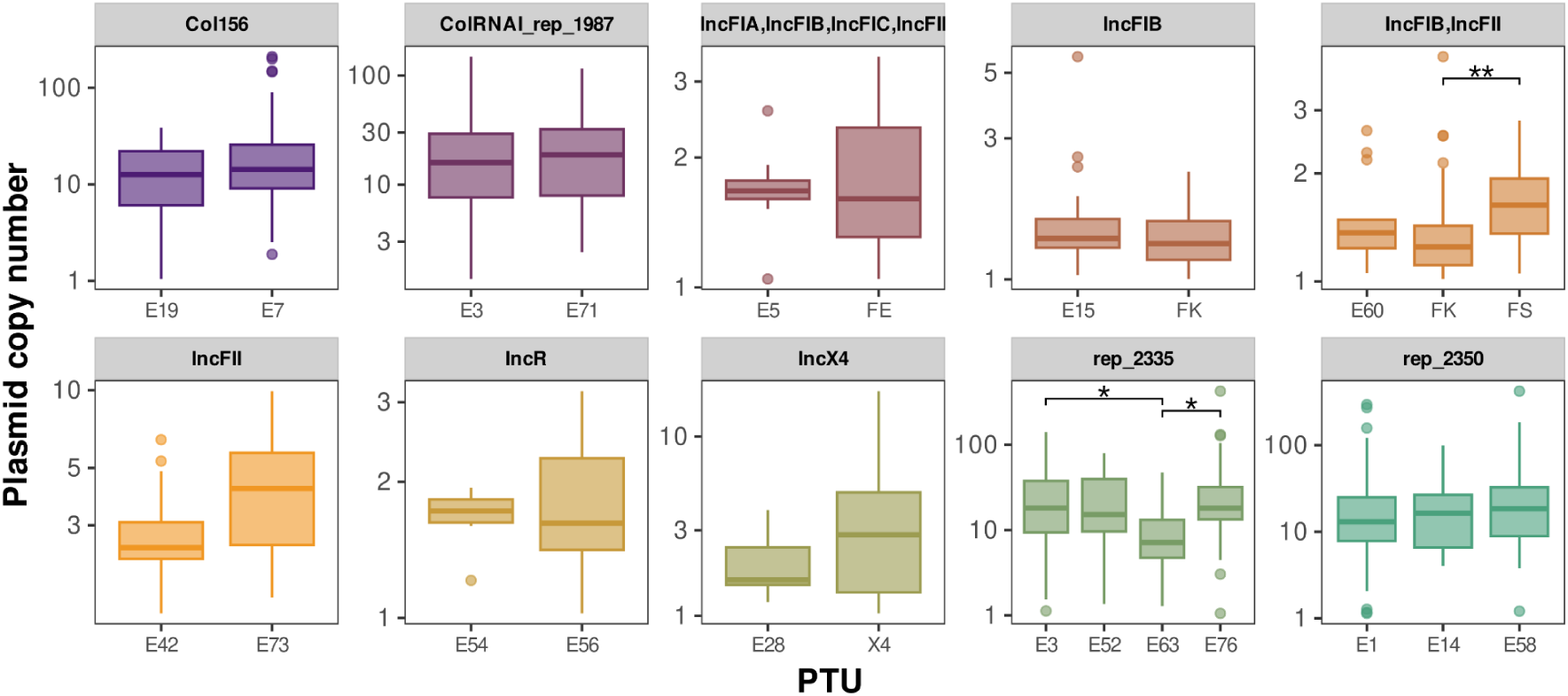
Plasmid copy number by PTUs within replicon types. Boxplots show the plasmid copy number (y-axis) across different PTUs (x-axis) for the most representative plasmid replicon types. Each panel represents a different plasmid replicon type (labeled at the top), with comparisons among its PTUs (labeled on the x-axis). Asterisks denote statistical significance according to Kruskal-Wallis test followed by Dunn’s test for pairwise multiple comparisons (* for p < 0.05, ** for p < 0.01).

**Supplementary Figure 6.**
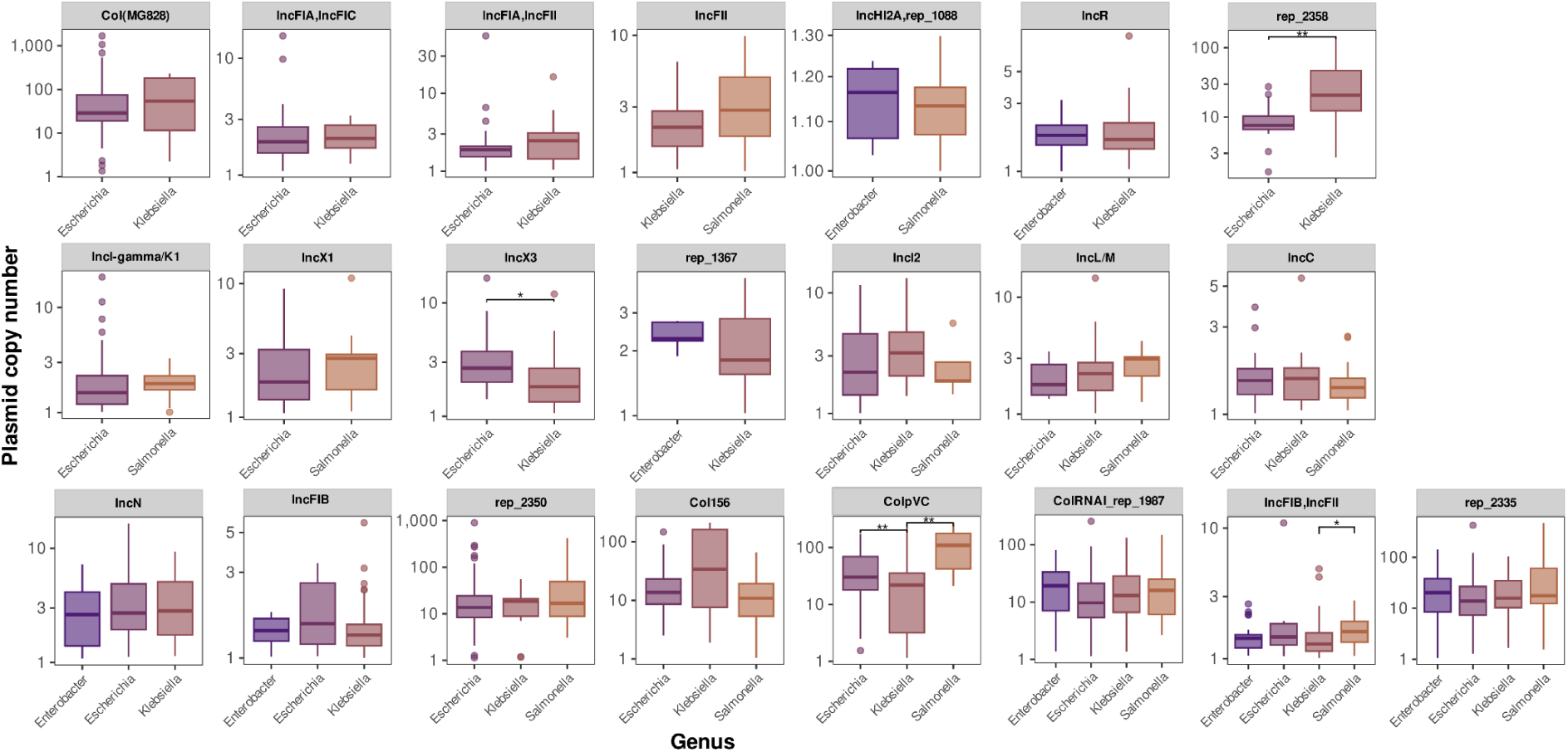
Plasmid copy number of replicon types across different hosts. Boxplots show the plasmid copy number (y-axis) across different hosts (x-axis) for the most representative replicon types in multiple genera. Each panel represents a replicon type (labelled at the top), with comparisons among host genera (labelled on the x-axis). Asterisks denote statistical significance according to the Kruskal-Wallis test, followed by Dunn’s test for pairwise multiple comparisons (* for p < 0.05, ** for p < 0.01).

**Supplementary Figure 7.**
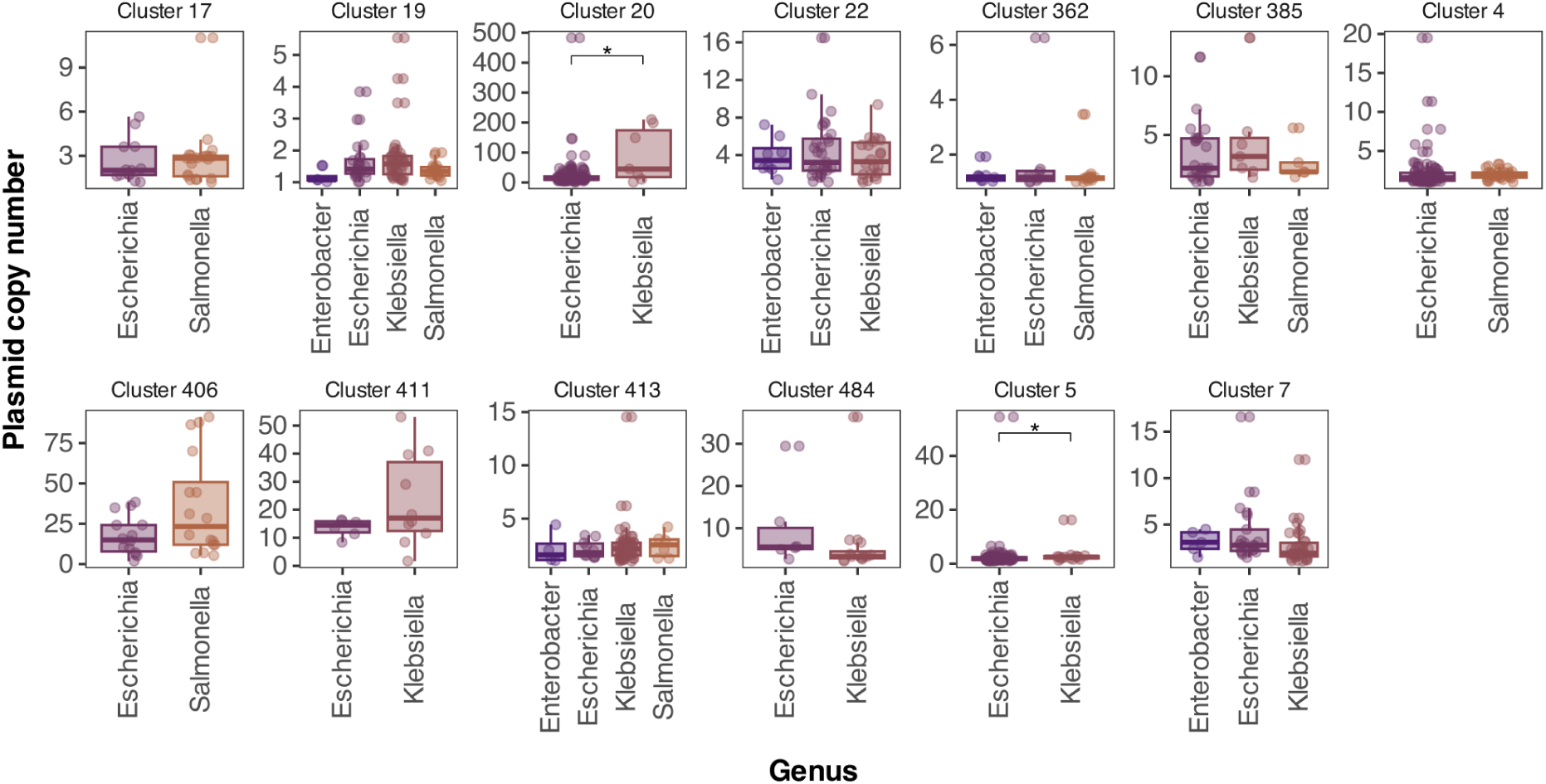
Plasmid copy number of plasmid clusters across different hosts. Boxplots show plasmid copy number (y-axis) distributions per host genus (x-axes) for the most representative clusters present in multiple genera (n > 3). Each panel represents a different plasmid cluster (labelled at the top), and each point represents a different plasmid. Asterisks denote statistical significance according to the Wilcoxon rank sum exact test, p < 0.047.

**Supplementary Figure 8.**
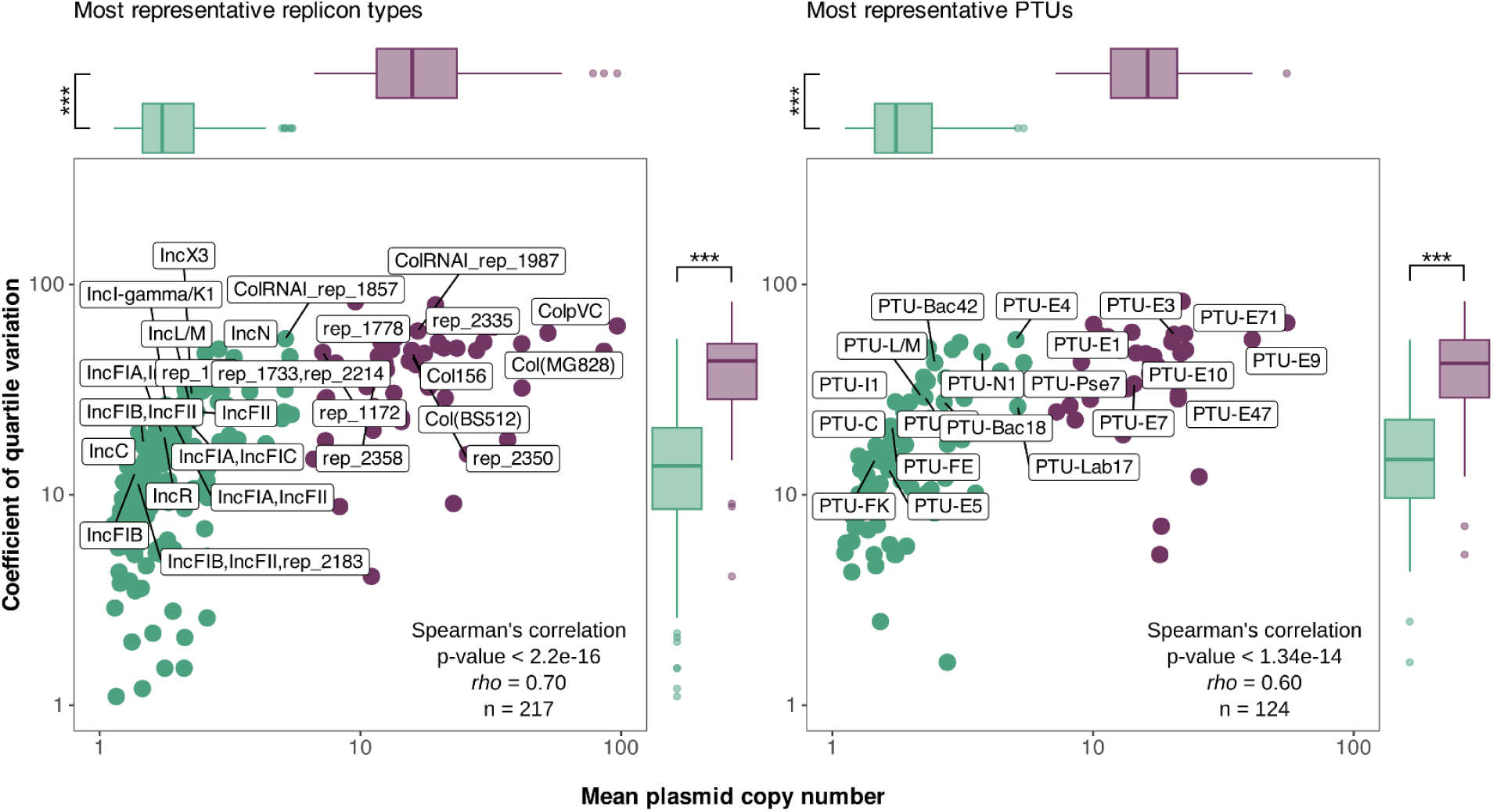
Intrinsic PCN variability, estimated without outliers. The mean PCN (x-axis) is plotted against the coefficient of quartile variation (CQV, y-axis), according to replicon type (left) or PTU (right), as in **Figure 3**. Each dot represents the mean PCN and CQV of a plasmid group, now estimated without outliers. The Spearman’s rho and p-value for each correlation are shown at the bottom right of each plot, and n indicates the sample size. Labels indicate the most representative plasmid groups. Colours distinguish HCP (purple) from LCP (green). Boxplots represent aggregated PCN (top) and CQV (right) data for all plasmid groups, according to the classification as HCPs and LCPs. Asterisks indicate significant differences between HCP and LCP (*** for p < 10^-7^, Wilcoxon rank sum exact test, effect size ≥ 0.404 for all tests of replicon types and ≥ 0.439 for all tests of PTUs), even after accounting by host shared ancestry (as estimated by Bayesian multilevel models with host as a random effect; see **Supplementary Dataset 6** for details).

**Supplementary Figure 9.**
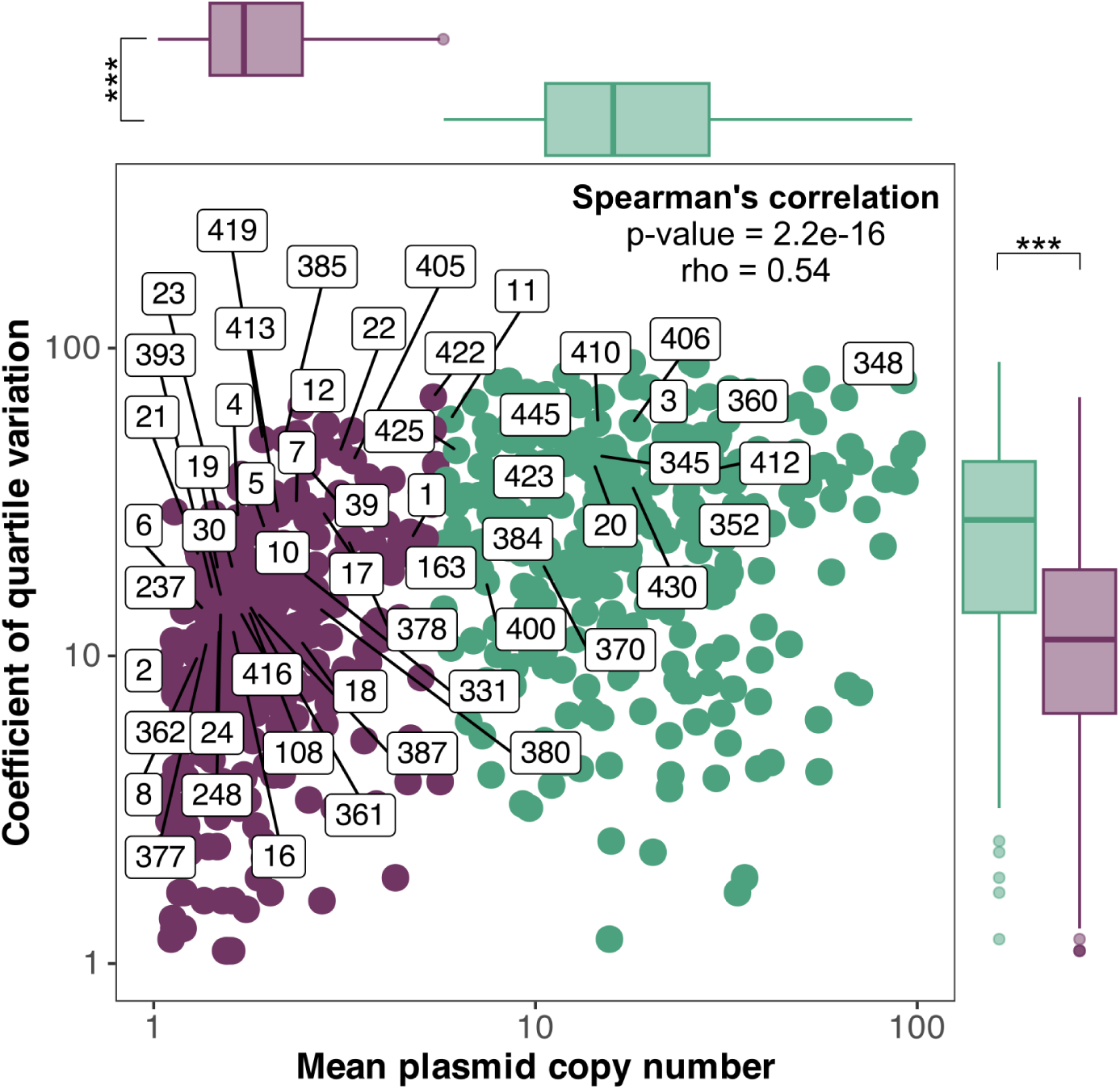
Intrinsic PCN variability per plasmid clusters. Each dot represents the mean PCN (x-axis) and CQV (y-axis) for the most representative plasmid clusters. Each cluster is labelled next to the respective point. The Spearman’s rho and p-value for the correlation are shown at the bottom right corner. Colours distinguish HCPs (purple) from LCPs (green). Boxplots represent aggregated PCN (top) and CQV (right) data for all plasmid groups, according to the classification as HCPs and LCPs. Asterisks indicate significant differences between HCP and LCP (*** for p < 0.001, effect size = 0.360; Wilcoxon rank sum exact test), even after accounting by host shared ancestry (as estimated by Bayesian multilevel models with host as a random effect; see **Supplementary Dataset 6** for details).

**Supplementary Figure 10.**
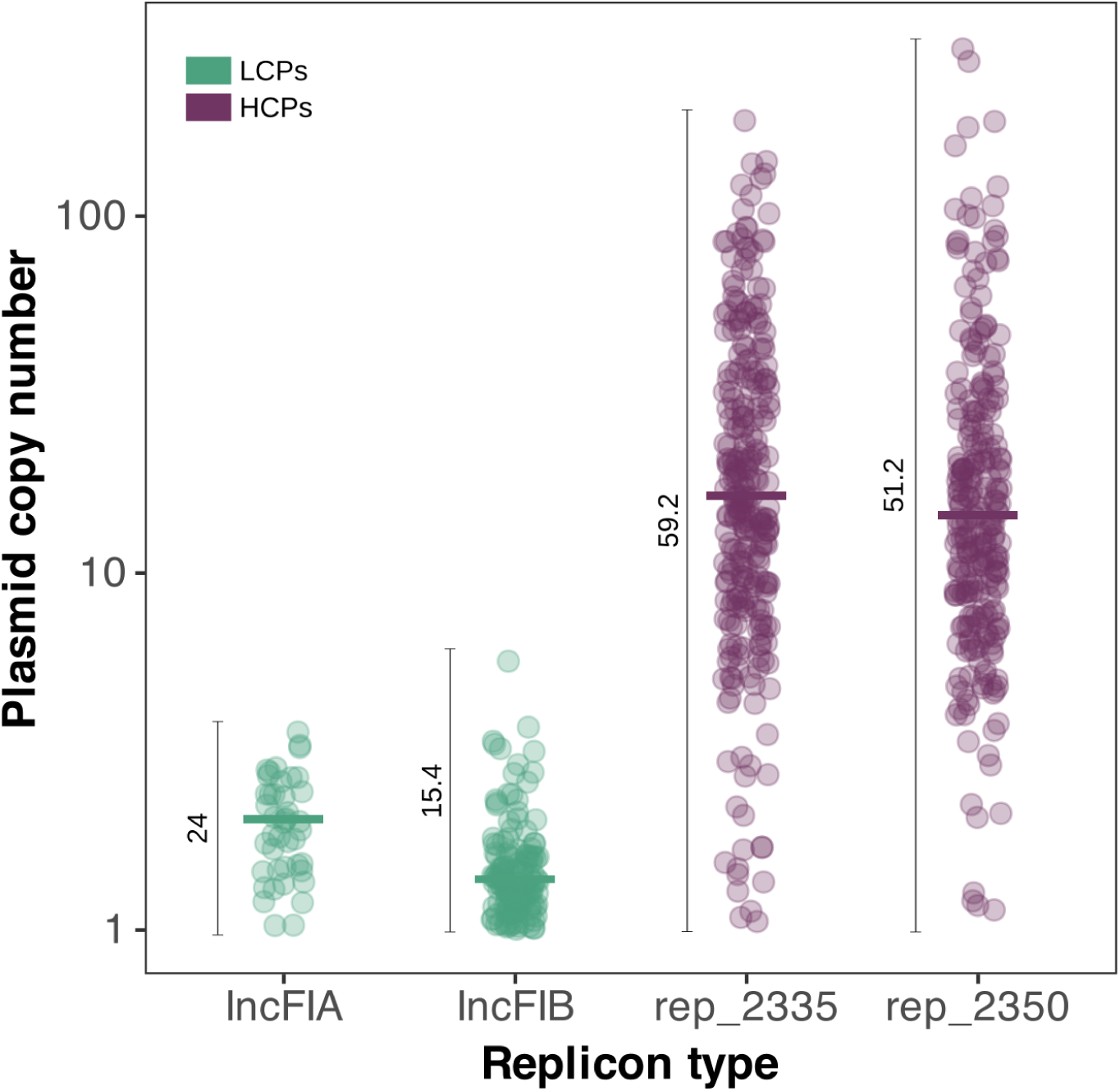
Intrinsic PCN variability of replicon types IncFIA, IncFIB, rep_2335, and rep_2350. The y-axis represents the plasmid copy number, and the x-axis represents replicon types. Colors represent low-copy plasmids (LCPs, green) and high-copy plasmids (HCPs, purple). Horizontal bars indicate the median plasmid copy number for each group. Vertical bars indicate the variability (CQV) in plasmid copy number for each replicon type.

**Supplementary Figure 11.**
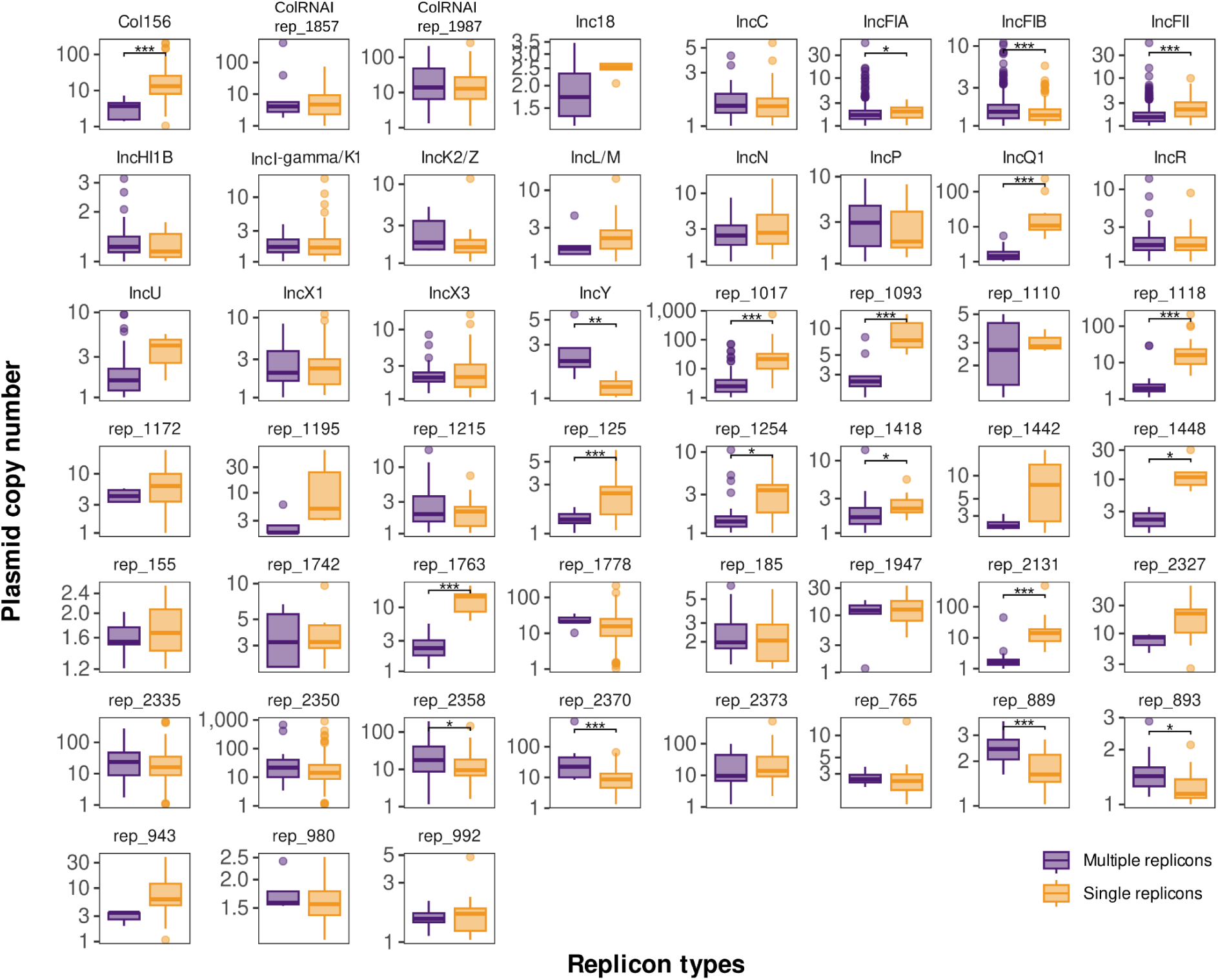
Variation in PCN between single and multi-replicon forms. The y-axis represents the plasmid copy number, and the x-axis represents the different plasmid forms: single replicons (orange) and multiple replicons (purple). Asterisks indicate statistically significant differences in PCN between single and multiple replicons (Wilcoxon rank sum exact test, * for p < 0.05, ** for p < 0.01, *** for p < 0.001).

**Supplementary Figure 12.**
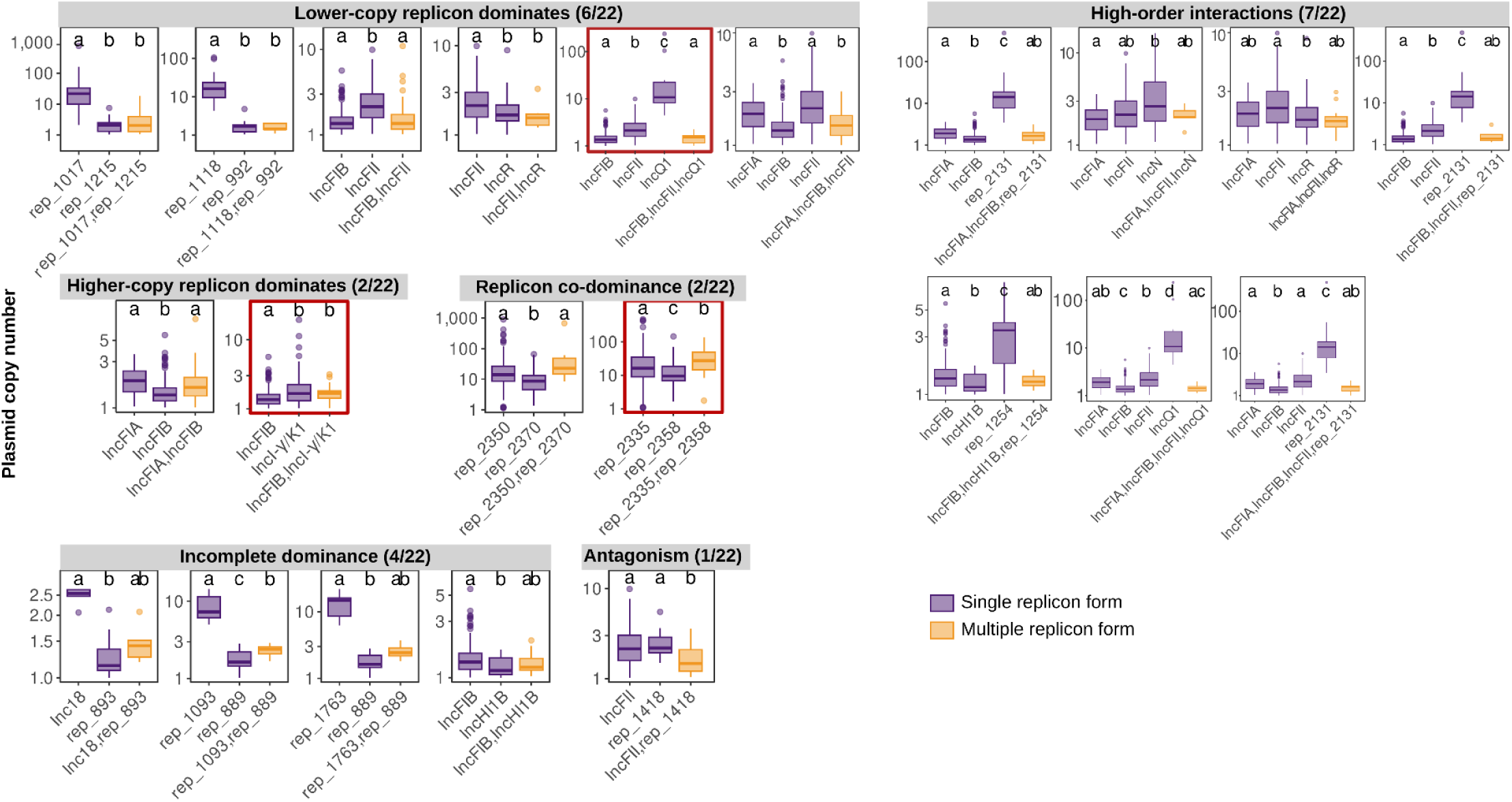
Replicon dominance interactions. Boxplots represent the plasmid copy number (y-axes) among different plasmid forms (x-axes): single replicons (purple) and multiple replicons (orange). Interactions are classified into six different groups: “Lower-copy replicon dominates”, “Higher-copy replicon dominates”, “Replicon co-dominance”, “Incomplete dominance”, “Antagonism”, and “High-order interactions. Letters assign groups based on statistical differences from the Dunn test (p < 0.05). Only groups with at least three samples are shown. Boxes highlighted in red correspond to the examples shown in **Figure 4 d-f**.

**Supplementary Figure 13.**
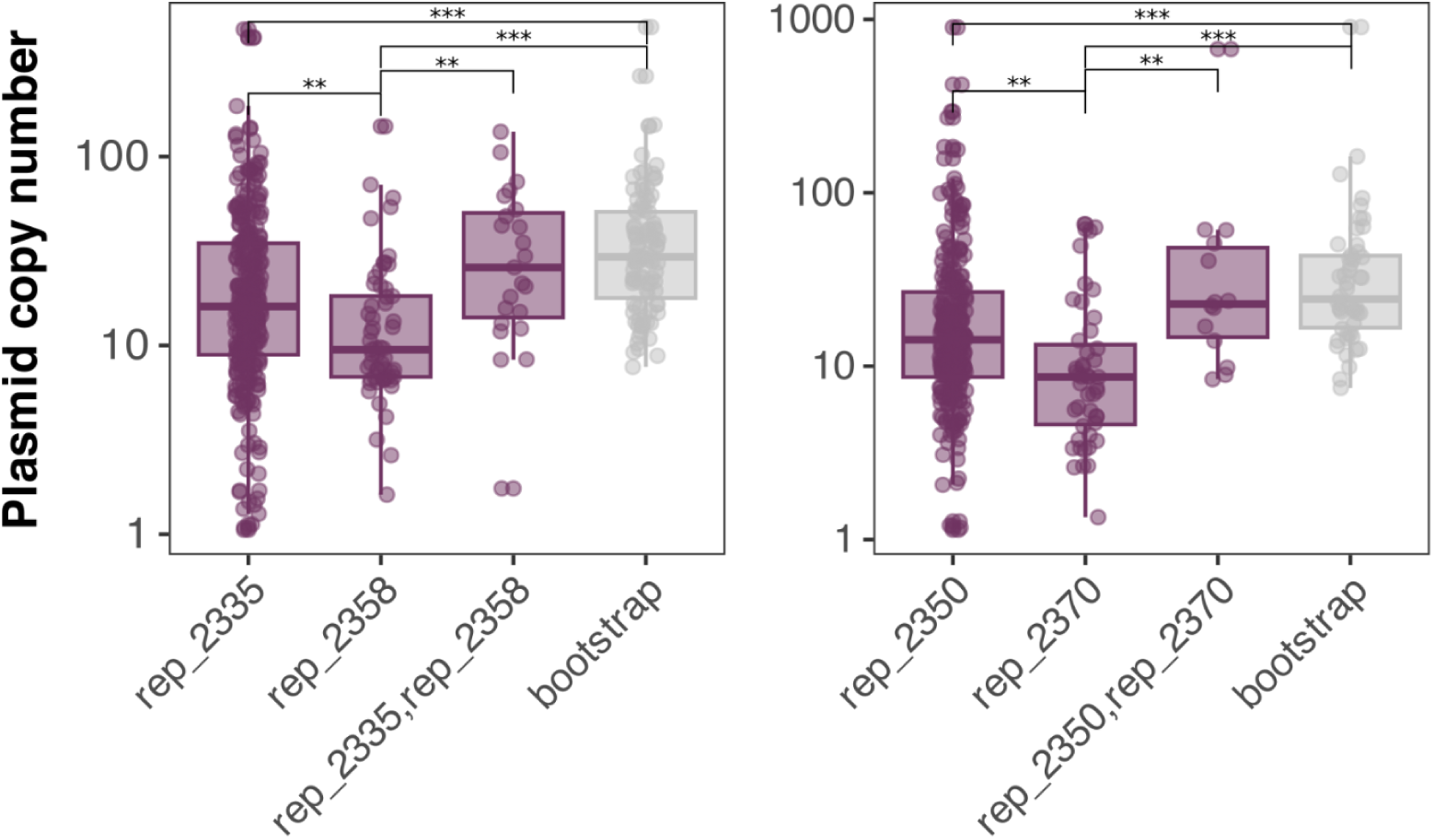
Candidate cases of additive effect in PCN due to replicon *co-dominance*. In each panel, boxplots represent the distribution of PCN (y-axis) for the different replicon states (x-axis). The observed PCN for each individual replicon and the multi-replicon form (rep_2335,rep_2358, and rep2350,rep2370) is indicated in purple. The expected PCN (additive), estimated through bootstrapping (see methods), is indicated in gray. Asterisks indicate statistically significant differences (Kruskal-Wallis test followed by Dunn’s test, ** p < 10^-2^, *** p < 10^-3^).

**Supplementary Figure 14.**
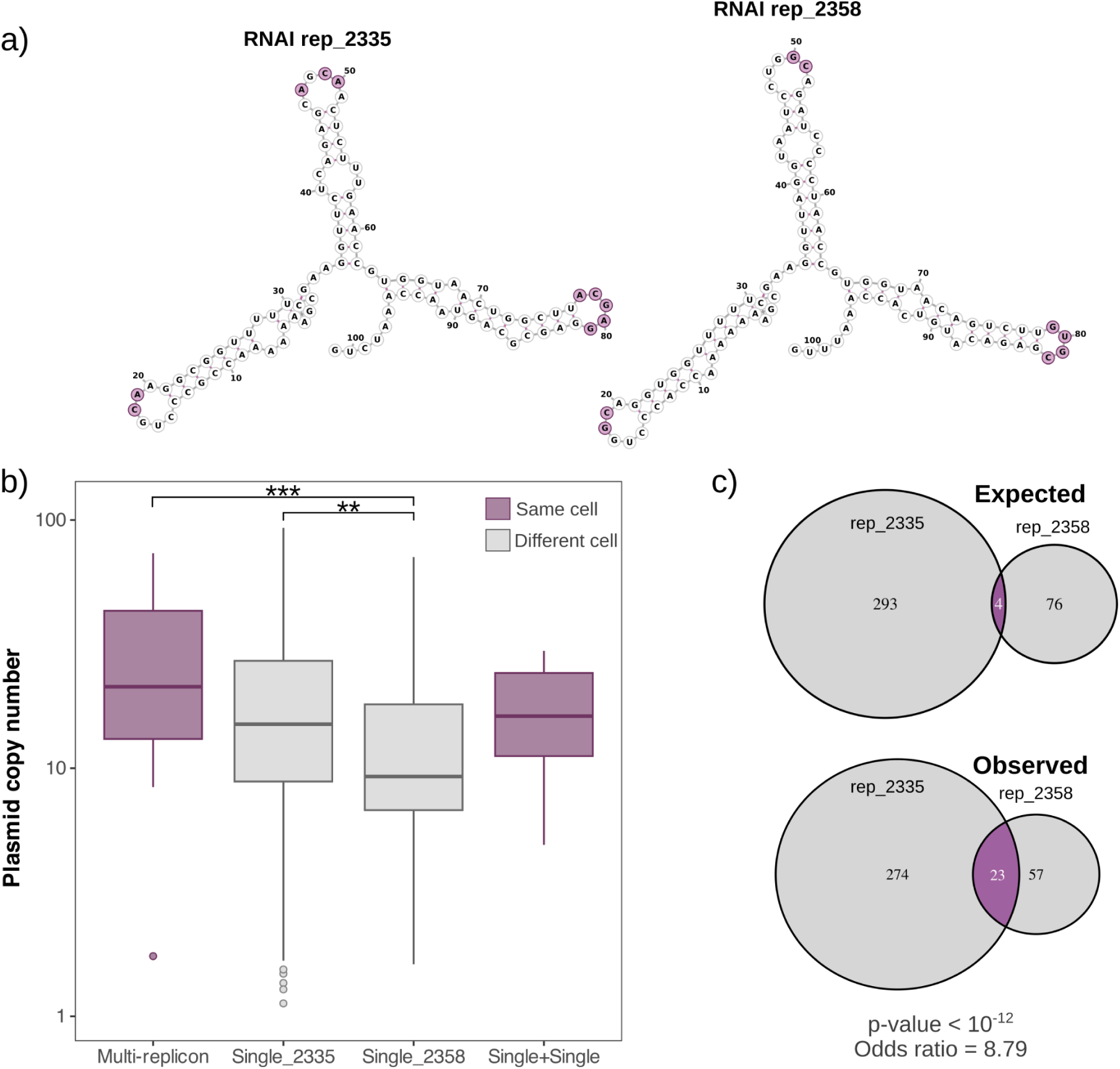
Structural analysis and PCN comparison of Col-like plasmids in different forms. **a)** Predicted RNAI structures from replicons rep_2335 and rep_2358 of the co-integrate example plasmid. RNAI negatively controls replication initiation in Col-like plasmids through three conserved stem-loops in its secondary structure. Therefore, mutations that affect these loops alter PCN and determine plasmid incompatibility. RNAI sequences were first extracted from the annotated plasmid sequences, and their secondary structures were predicted using the RNAfold Webserver (http://rna.tbi.univie.ac.at//cgi-bin/RNAWebSuite/RNAfold.cgi). **b)** The boxplots show the PCN of two replicons when they are found individually in different cells (“Single_2335”, “Single_2358”), when they are co-integrated in the cell (“Multi-replicon”) or co-residing as individual replicons within the same cell (“Single+Single”) (Kruskal-Wallis test followed by Dunn’s test for pairwise multiple comparisons, ** p < 10^-2^, *** p < 10^-3^), indicating that these two replicons were likely compatible and that their combined PCN was additive. **c)** The Venn diagrams show expected and observed counts for rep_2335, rep_2358, and genomes carrying co-existing single-replicon plasmids. Genomes bearing both single-replicon were overrepresented (Fisher’s exact test p < 10⁻¹², odds ratio = 8.79), indicating that these two replicons are likely compatible.

**Supplementary Figure 15.**
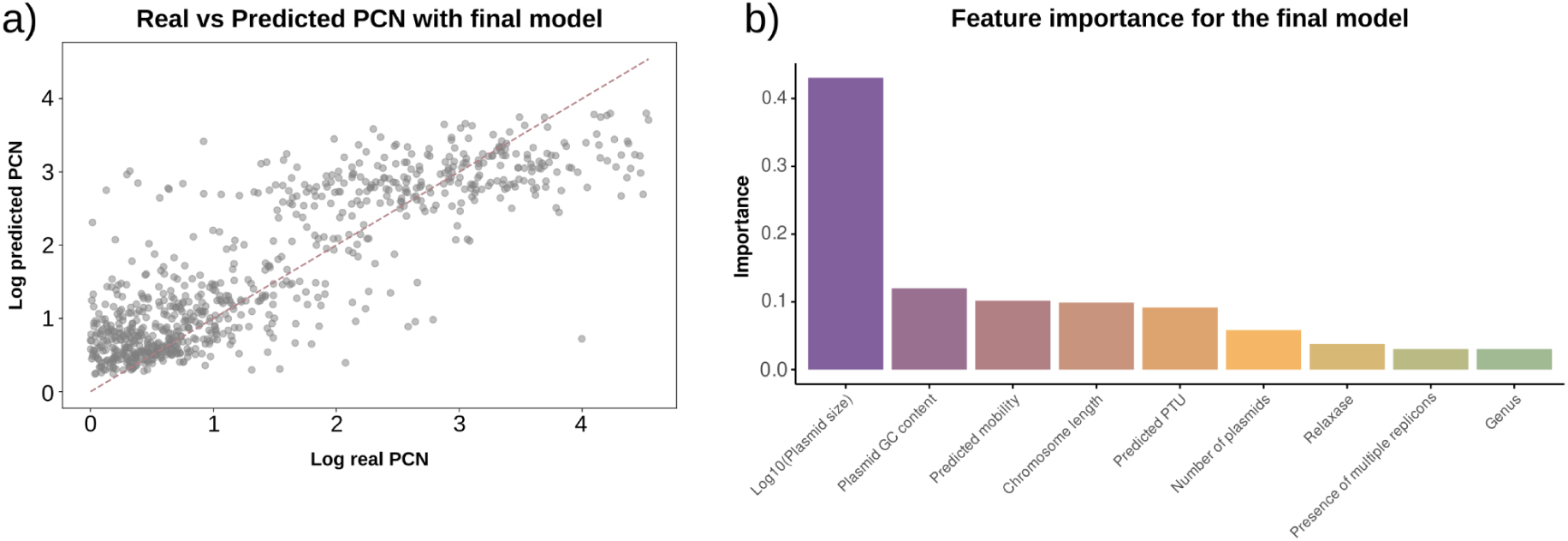
Analysis of factors influencing plasmid copy number (PCN). **a)** Performance of the Random Forest Regressor model in predicting PCN. The scatter plot shows predicted vs. observed PCN values. Each point represents an individual plasmid. **b)** Gini Feature importance plot from the Random Forest Regressor model. Bar height represents the relative importance of each variable for predicting PCN.

**Supplementary Figure 16.**
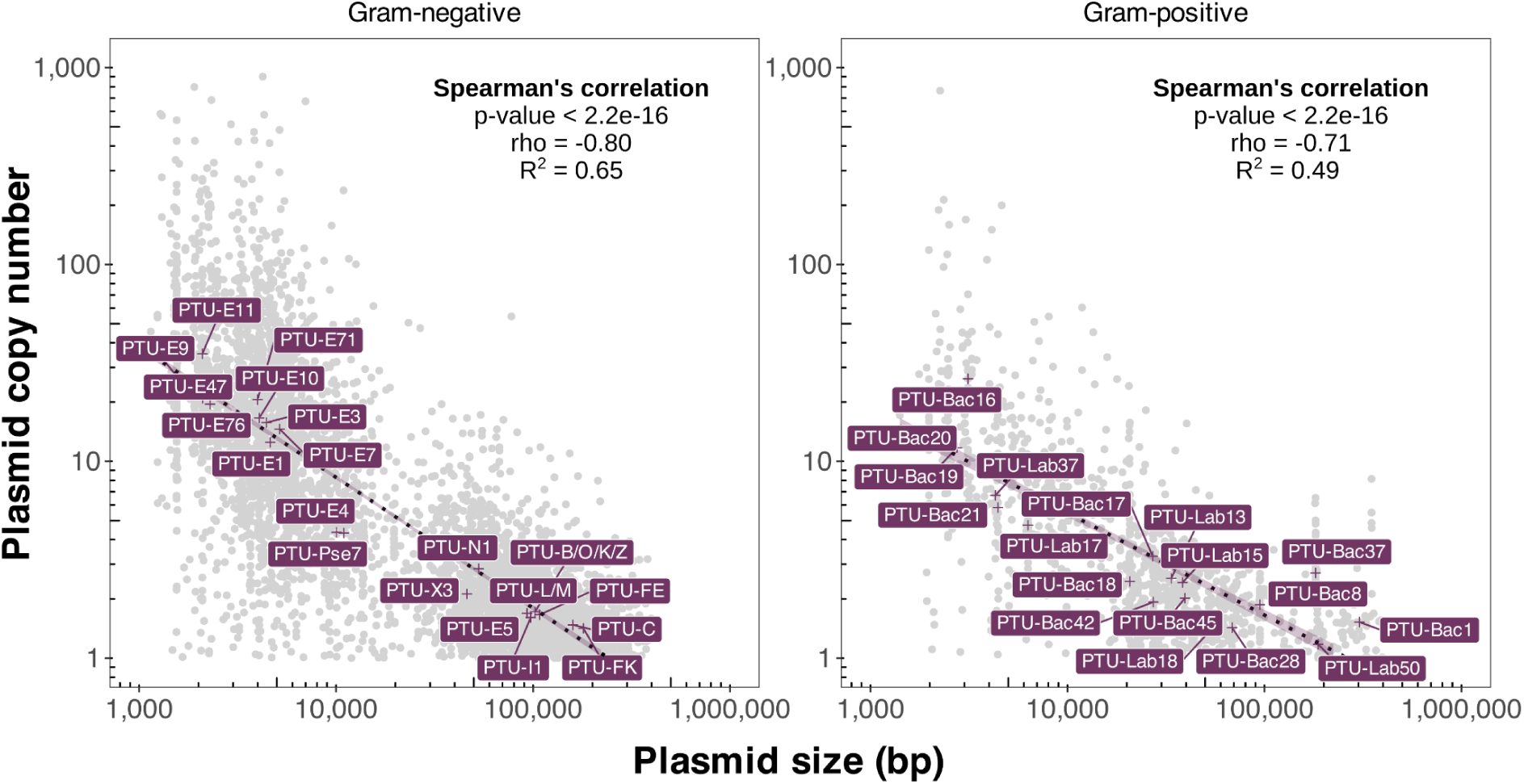
The negative correlation between plasmid size and copy number is conserved. Correlations between PCN (y-axis) and plasmid size (x-axis) for both Gram-positive and Gram-negative bacteria. Each dot represents a single plasmid, and the dotted line represents the best linear fit with 95% confidence intervals in light purple. The purple crosses and labels indicate each PTU’s median PCN and size. Spearman’s p-value and rho for correlations, and R^2^ of the linear model (**Supplementary dataset 7**) are also indicated within each plot (n = 5,181 for Gram-negative and n = 1,146 for Gram-positive).

**Supplementary Figure 17.**
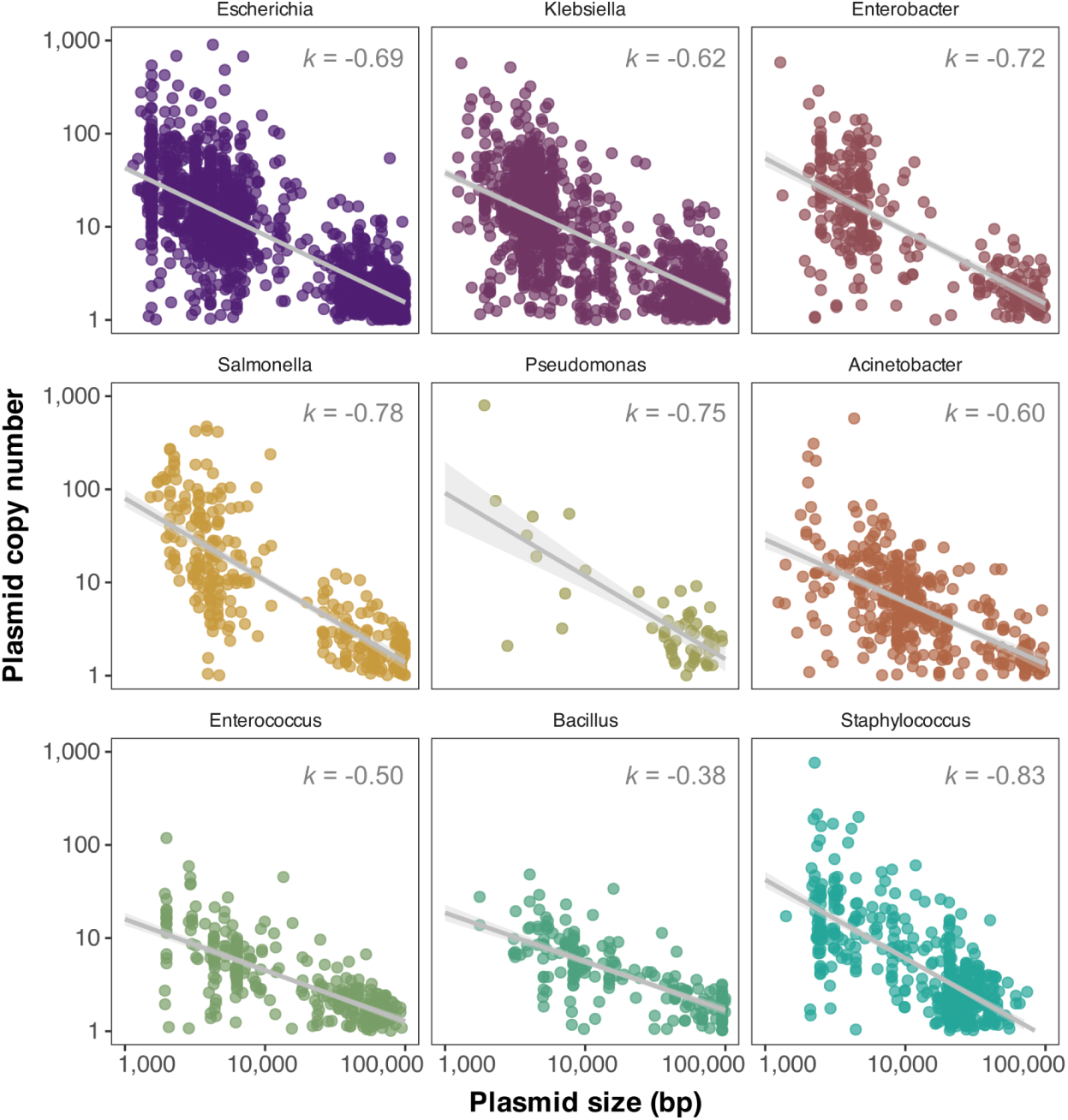
Relationship between plasmid size and copy number across bacterial genera. The scatter plots show the correlation between plasmid size (x-axis) and plasmid copy number (y-axis) for each genus. Each point represents an individual plasmid. The correlation is displayed by an ordinary least squares regression line (in gray), with the surrounding shaded area indicating the 95% confidence interval. The slope of the regression (*k*) is annotated at the top-right corner of each plot.

**Supplementary Figure 18.**
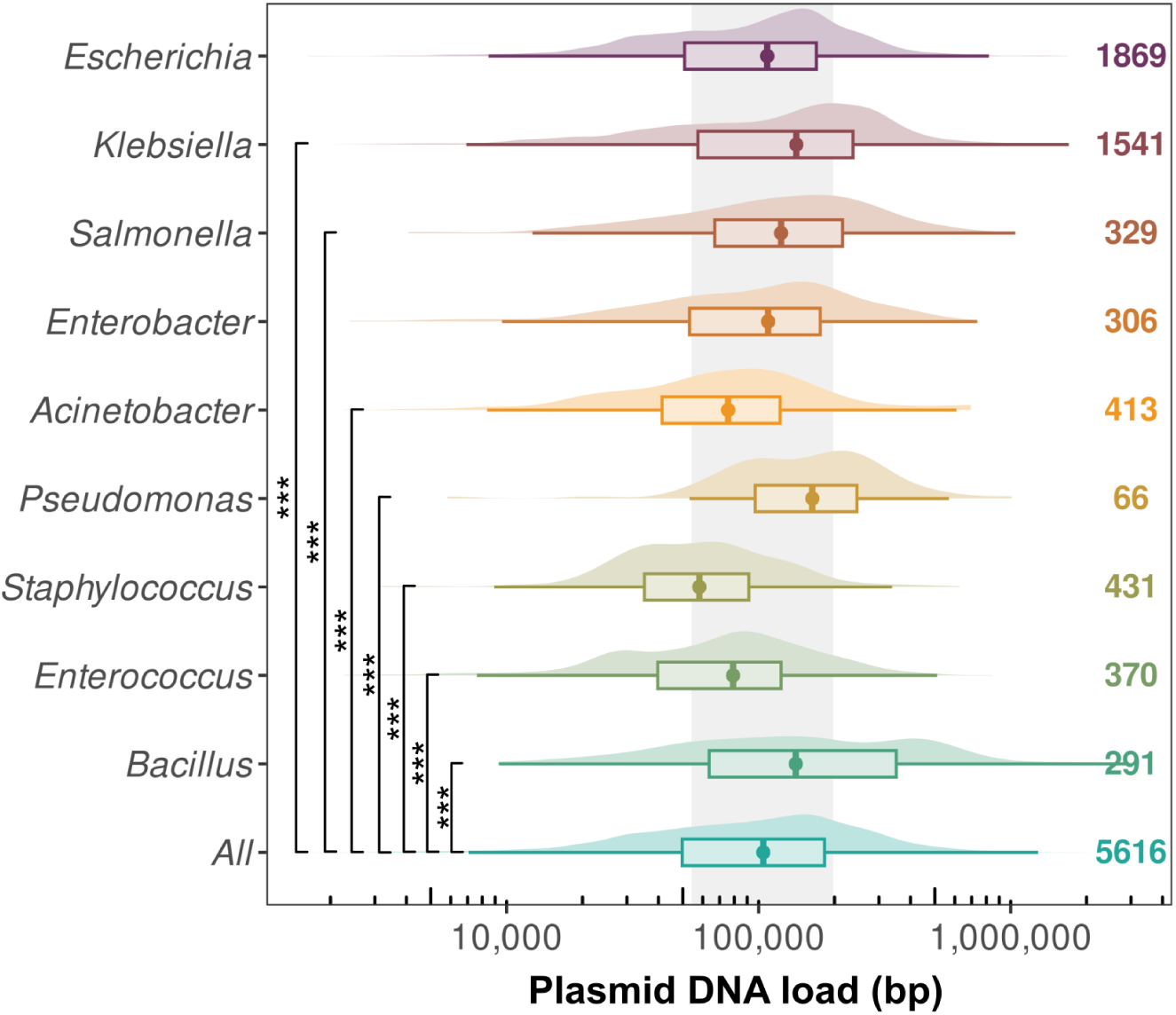
Distribution of total plasmid DNA across different bacterial genera. The y-axis represents the analyzed genera, while the x-axis represents the total DNA load per plasmid (bp), calculated as the product between PCN and plasmid size. The distributions (with boxplots overlaid) represent total plasmid DNA per genus (Kruskal-Wallis test followed by Dunn’s test for pairwise multiple comparisons, p-value < 10^-4^).

**Supplementary Figure 19.**
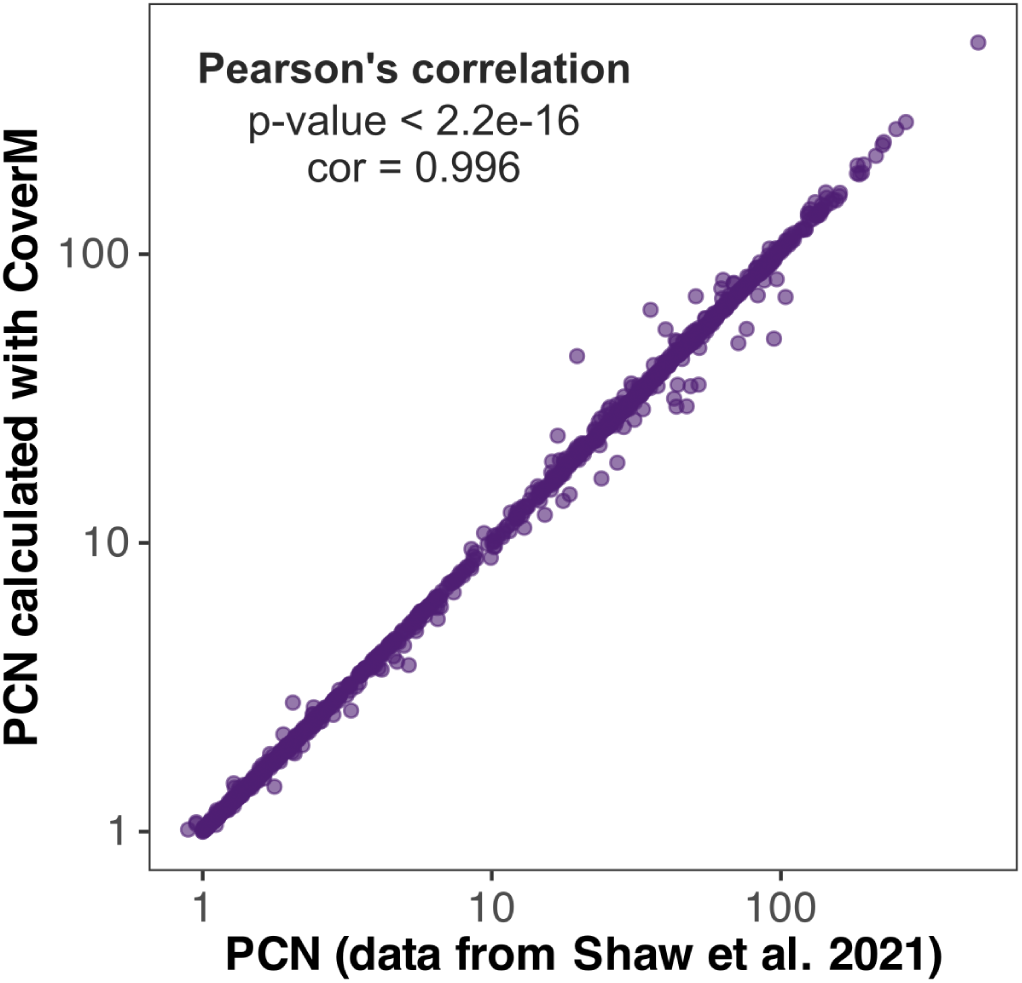
Correlation of plasmid copy number estimates between methods. The x-axis represents PCN data calculated from Illumina sequencing from reference^17^, and the y-axis represents PCN calculated using CoverM. Each point represents an individual plasmid. Pearson’s correlation coefficient (r = 0.996) and the associated p-value (< 10^-16^) are shown in the top-left corner.

**Supplementary Figure 20.**
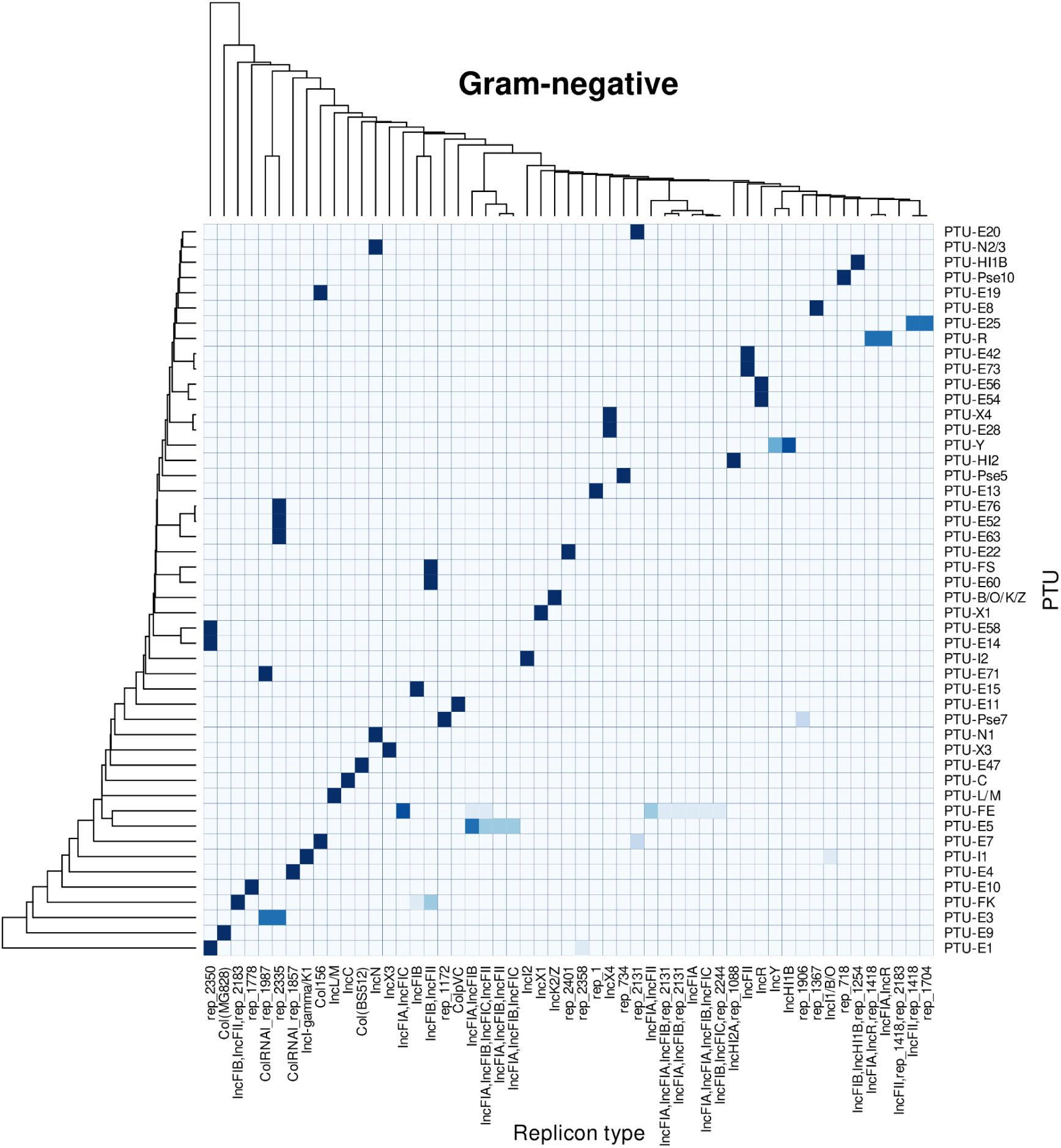
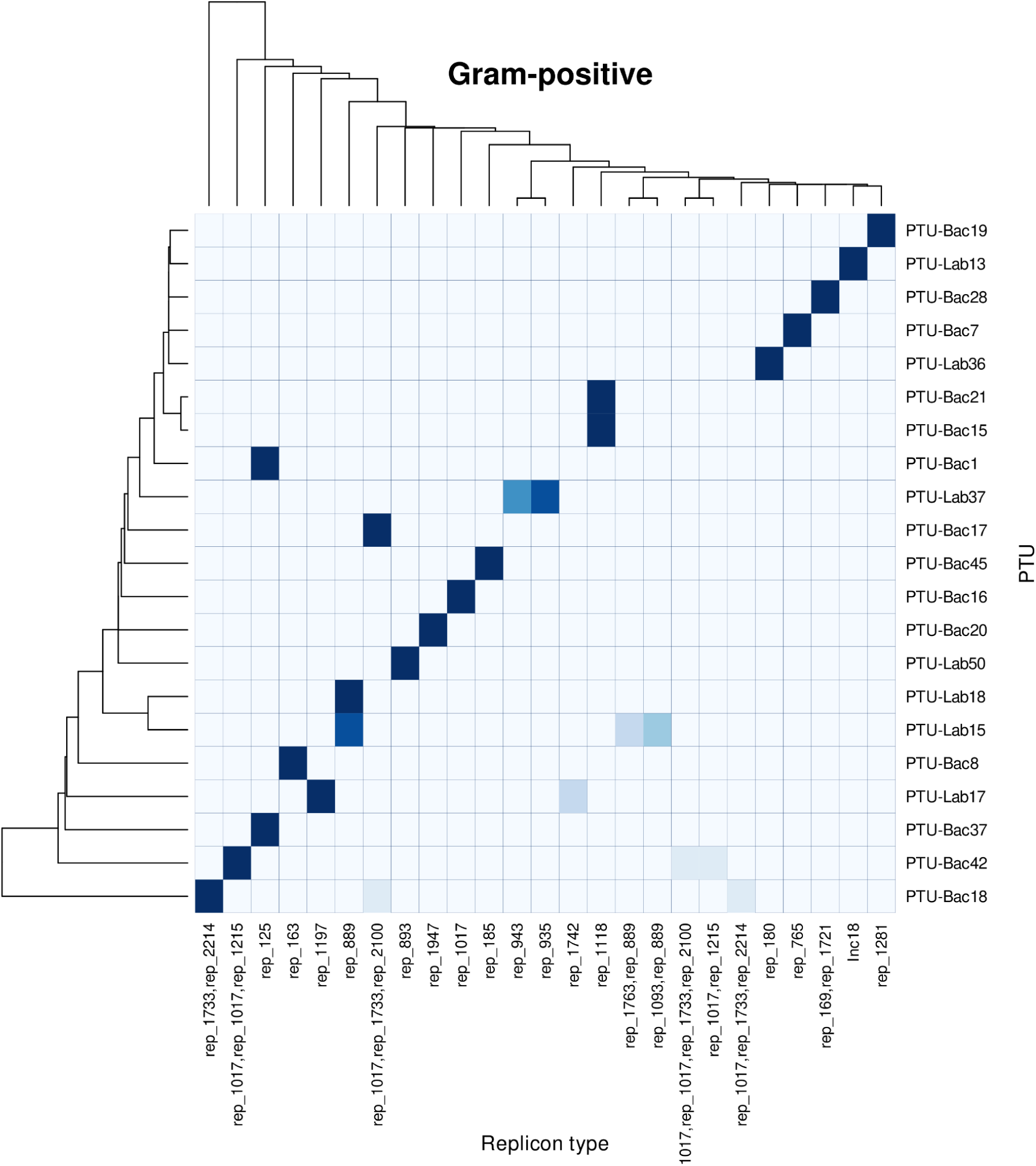
Relation between PTUs and replicon types in Gram-negative hosts. Heatmap displaying the association between PTUs (y-axis) and replicon types (x-axis) in Gram-negative bacteria. Based on their similarity, the dendrograms on the left and top of the heatmap show the hierarchical clustering of PTUs and replicon types. Each cell of the heatmap represents the strength of the association between a PTU and a replicon type. Darker shades of blue signify a stronger association, while lighter shades indicate weak or nonexistent associations. Groups with an occurrence of <10 for Gram-negative and <5 for Gram-positive bacteria are not displayed.

